# Repeatability of protein structural evolution following convergent gene fusions

**DOI:** 10.1101/2025.02.23.639786

**Authors:** Naoki Konno, Keita Miyake, Satoshi Nishino, Kimiho Omae, Haruaki Yanagisawa, Saburo Tsuru, Yuki Nishimura, Masahide Kikkawa, Chikara Furusawa, Wataru Iwasaki

## Abstract

Convergent evolution of proteins provides insights into repeatability of genetic adaptation. While local convergence of proteins at residue or domain level has been characterized, global structural convergence by inter-domain/molecular interactions remains largely unknown. Here we present structural convergent evolution on fusion enzymes of aldehyde dehydrogenases (ALDHs) and alcohol dehydrogenases (ADHs). We discovered BdhE (bifunctional dehydrogenase E), an enzyme clade that emerged independently from the previously known AdhE family through distinct gene fusion events. AdhE and BdhE showed shared enzymatic activities and non-overlapping phylogenetic distribution, suggesting common functions in different species. Cryo-electron microscopy revealed BdhEs form donut-like homotetramers, contrasting AdhE’s helical homopolymers. Intriguingly, despite distinct quaternary structures and >70% unshared amino acids, both enzymes form resembled dimeric structure units by ALDH-ADH interactions via convergently elongated loop structures. These findings suggest convergent gene fusions recurrently led to substrate channeling evolution to enhance two-step reaction efficiency. Our study unveils structural convergence at inter-domain/molecular level, expanding our knowledges on patterns behind molecular evolution exploring protein structural universe.

## MAIN TEXT

Repeatability is fundamental to science, distinguishing non-random from random phenomena and elucidating causality and predictability (*1–3*). Evolution in nature, however, poses challenges for studying repeatability, since we can typically examine historical events that have occurred just once (*4*, *5*).

Evolution is characterized by stochastic mechanisms (e.g., mutations and genetic drift), and the adaptation processes are heavily influenced by historical contingencies (*4*, *5*). These effects decrease the likelihood of repeated evolution, even when subjected to similar selective pressures. Convergent evolution— independent yet similar evolutionary outcomes in distinct lineages—therefore, is a crucial key to discuss evolutionary repeatability.

Through molecular evolution studies, convergence of proteins has been characterized at various levels to decipher driving factors (e.g., selective pressures and evolutionary constraints) and predictability of genetic adaptation (*6–8*). At residue level, different lineages of a protein family show independent but same mutations at specific residues under similar selective pressures, such as mutations of ATPα conferring plant toxin resistance to insects (*9–11*). At function level, some evolutionarily unrelated enzyme families have evolved similar catalytic activity by acquiring similar active sites, like Ser-His-Asp triad of trypsin and subtilisin (*8*, *12*). Other studies have unveiled structural convergence at domain level (e.g., Cren7/Sul7 and SH3 domains) (*13*, *14*). While all these previous studies revealed convergent evolution at local parts of a protein molecule (i.e. specific residues or domains), convergence of more global three dimensional (3D) structures (i.e., whole structures of protein monomers or multimers) by recurrent evolution of similar inter-domain and inter-molecular interactions remains largely unexplored.

Such protein global structure evolution can be triggered by drastic genetic changes such as gene fusions and fissions (*15*, *16*). These events alter domain architectures of proteins, and especially gene fusion events can further lead to fixation of spatial proximity between previously uncontacted protein domains via linkers (*17–19*). The fixed proximity between domains let them collide and contact each other frequently and potentially facilitates global protein structure evolution by acquiring inter-domain interactions. Although remarkable number of domain architecture convergence have been suggested by bioinformatic analyses (*20*, *21*), it is poorly understood whether recurrent 3D structural evolution follows repeated domain architecture emergence.

Alcohol/aldehyde dehydrogenases, comprising aldehyde dehydrogenase (ALDH) and alcohol dehydrogenase (ADH) domains, present suitable targets for studying structural evolution driven by gene fusions. While most eukaryotes, including humans, possess separate genes encoding ALDH and ADH domains, some bacterial species are known to harbor AdhE, a previously known protein family consisting of both ALDH and ADH domains. The ALDH and ADH domains of AdhE are most closely related to EutE and PduQ families, respectively (*22*, *23*), and are thought to have emerged through gene fusions of the ancestral single-domain ALDH and ADH genes. With this unique domain composition, AdhE forms helical filament structures as a homopolymer, termed spirosomes initially reported in 1970s (*24*). In the recently solved structure of spirosomes, adjacent AdhE molecules exhibit inter-molecular interactions between ALDH and ADH domains (*25–28*). This interaction was reported to facilitate substrate channeling between ALDH and ADH domains, passing aldehydes as reaction intermediates (*25*, *26*). This structural arrangement is thereby considered adaptive, because it enables efficient catalysis of the two serial reactions, simultaneously preventing the leakage and diffusion of the cytotoxic aldehydes in cells (*25*, *26*). Furthermore, the pitch of the helical structure has been found to change like a spring triggered by cofactor bindings, proposed as a mechanism of enzyme activity regulation (*26*, *28*).

In this study, we first report a discovery of another bifunctional aldehyde/alcohol dehydrogenase (BdhE) that evolved through a gene fusion of ALDH and ADH genes, similarly yet independently to AdhE. Although AdhE and BdhE unshared more than 70% amino acid residues, we demonstrated that BdhE shares functional characteristics with AdhE. We further solved and compared the 3D structures of recombinant AdhE and BdhE, and unveiled distinct multimer conformations yet similar inter-domain and inter-molecular interactions between AdhE and BdhE, illuminating the extent of repeated structural evolution after independent gene fusions. We further dug into pre-existing genomic contexts of the convergent gene fusions to decipher how similar fusion enzymes repeatedly evolved in bacteria. Our work broadens the current understanding of repeatability in molecular evolution and suggests potentially universal evolutionary patterns for 3D structural evolution of proteins.

### BdhE is an ALDH-ADH protein family but is evolutionarily distinct from AdhE

Through a sequence homology search for *adhE* across 45,555 bacterial representative genomes from the Genome Taxonomy Database (GTDB) (*29*), we serendipitously identified a set of genes exhibiting low sequence identity (<30%) but high alignment coverage (>90%) with known *adhE* genes (**Fig. 1A**). These genes (temporarily named quasi-*adhE*) coded proteins with predicted structures like that of AdhE (**Fig. 1B**). Notably, the sequence identities were even lower than those partially aligned with AdhEs (40-50% coverage), including single-domain proteins containing only ALDH or ADH (**Fig. 1A**). Based on these observations, we hypothesized that at least one protein family originated by fusions of ALDH and ADH genes and independently from the fusion event that gave rise to AdhE.

**Figure 1.**
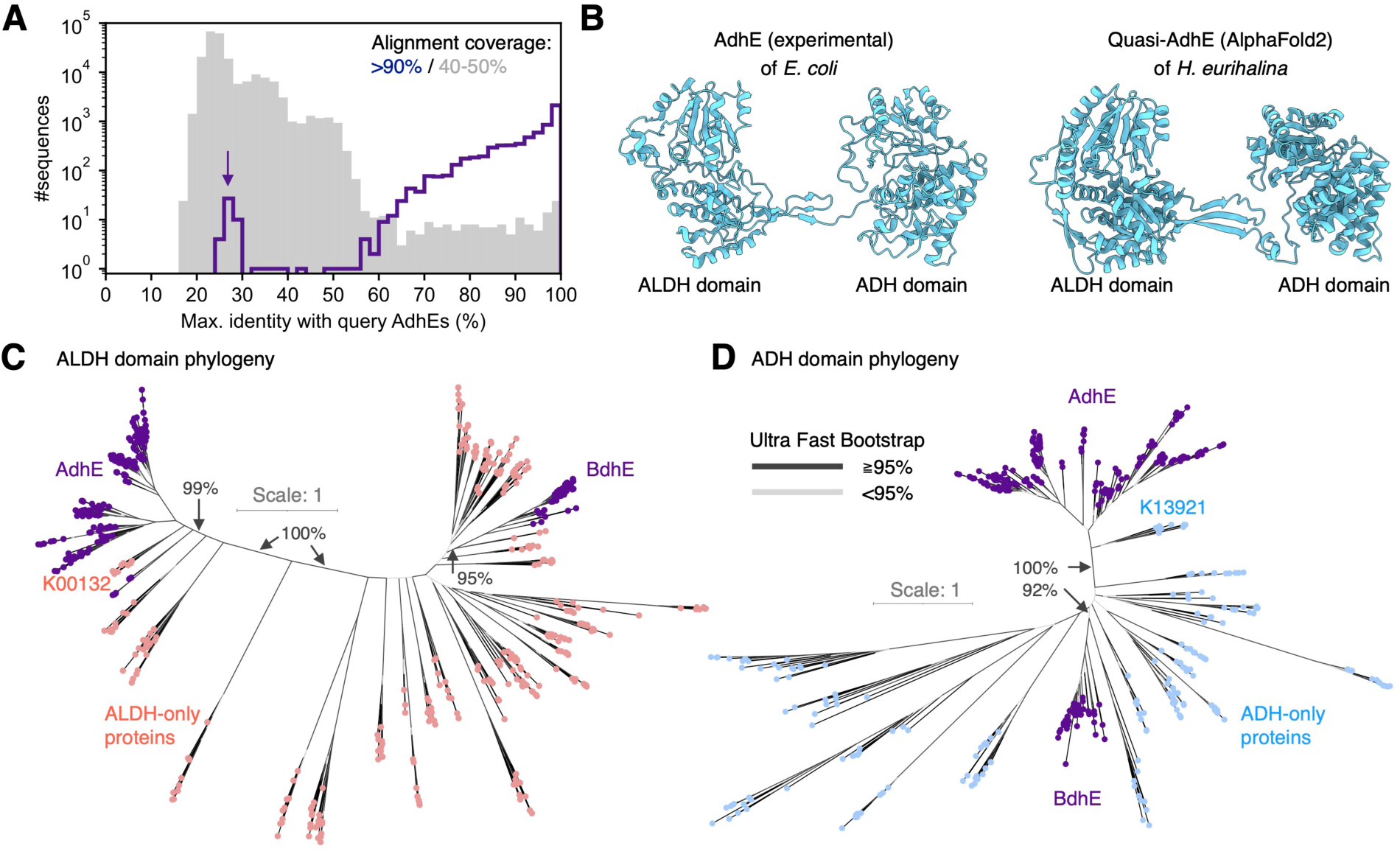
**AdhE and BdhE possessing similar yet evolutionarily independent domain architectures**. **(A)** Histogram of maximum alignment identities (%) of bacterial proteins hit by sequence similarity search querying known AdhE sequences. The purple line and the grey shade represent the histograms for hit proteins showing >90% and 40-50% alignment coverages, respectively. The arrow indicates the peak of genes showing high alignment coverages but low identities with AdhE (namely Quasi-AdhE). **(B)** An experimentally determined monomer structure of AdhE (compact form; PDB ID: 6TQM) and a predicted monomer structure of Quasi-AdhE from *Halomonas eurihalina* by AlphaFold v2.3.2. **(C, D)** Gene phylogenies of proteins possessing ALDH (C) or ADH (D) domain. We conducted multiple sequence alignment and phylogeny estimation for the union of 200 randomly sampled AdhEs and all 47 BdhEs (Quasi-AdhEs), as well as 10 proteins randomly selected for every KEGG Ortholog only with ALDH or ADH domains. The red, blue, and purple tips represent ALDH-only, ADH-only, and ALDH-ADH fusion proteins, respectively. The black and grey branches represent splits for which the Ultrafast Bootstrap values were ≧95% or not, respectively.

To comprehensively identify proteins with the same domain composition as AdhE, we retrieved all 8,467 proteins with ALDH-ADH composition from InterPro database (*30*). Notably, only five proteins in InterPro exhibited the reversed ADH-ALDH composition. Ortholog assignment for the ALDH-ADH proteins revealed 44 proteins not annotated as AdhE. Of these, 23 proteins exceeded 800 amino acids in length and exhibited low association with the AdhE family (**fig. S1A, B**). These findings suggest that these 23 proteins likely belong to Quasi-AdhE families. We used these Quasi-AdhEs as queries to search for homologous genes in 45,555 GTDB genomes (**fig. S1C, D**). Then, we reconstructed gene phylogenies for both ALDH and ADH domains of the search hit genes, rooted with an AdhE sequence from *Escherichia coli* (*E. coli*). Genes with ALDH and ADH domains formed a monophyletic clade apart from the outgroup AdhE in both phylogenies, supported by 100% Ultrafast Bootstrap (UFboot) values (**fig. S2**). We thereby named this protein family BdhE (“Bifunctional dehydrogenase E”).

To elucidate the evolutionary relationship between AdhE and BdhE, we estimated phylogenetic trees across orthologs with ALDH and/or ADH domain, including AdhEs and BdhEs (**Fig. 1C, D**). The results revealed that both ALDH and ADH domains of AdhE and BdhE formed distinct clades separated by multiple branches with 99% or more UFboot values. The sister groups of AdhE’s ALDH and ADH domains were “acetaldehyde dehydrogenase (acetylating)” (K00132 in KEGG Orthology; including genes annotated as “EutE” consistently with a previous report (*22*)) and “1-propanol dehydrogenase, *pduQ*” (K13921), supported by 99% and 100% UFboot values, respectively. Although the sister groups of BdhE were not supported by sufficiently large UFboot values (>95%) in **Fig. 1C** and **D**, phylogenies of proteins closest to BdhE indicated “aldehyde dehydrogenase” (K00128) and “maleyl acetate reductase” (K00217) as sister clades of BdhE with 99% and 100% UFboot values (**fig. S2**). These results suggest that while AdhE and BdhE share the same domain architecture, they originated from distinct fusion events of evolutionarily distant ALDH and ADH genes. Consequently, AdhE and BdhE present a suitable model case for studying convergent gene fusions and subsequent functional and structural evolution.

### BdhE showed shared enzymatic activity and non-overlapped phylogenetic distribution with AdhE

To elucidate the enzymatic function of BdhE, for which no activity had been experimentally reported or registered in UniProt, we expressed and purified BdhE from *Halomonas eurihalina* and the previously characterized AdhE from *E. coli* (*25*, *26*) under the same conditions (**fig. S3A**). We assayed their activities for ethanol oxidation and acetyl-CoA reduction adjusting to previously reported assay conditions for AdhE (*25*). The absorbance at 340 nm, indicative of nicotinamide adenine dinucleotide (NADH) concentration, changed remarkably upon the addition of either AdhE or BdhE compared to the no enzyme control although BdhE showed lower reaction rates than AdhE (**Fig. 2A, B, fig. S3B and C**). We also confirmed aldehyde reduction activity of ADH domain (**fig. S3B**). These results align with reports of reversible ethanol-acetyl-CoA conversion of AdhE (*25*, *26*, *31*). We further tested choline oxidization activity as BdhE is evolutionarily close to betaine aldehyde dehydrogenase (K00130) (**fig. S2**), then found that both enzymes showed the activity (**fig. S3C**). These results consistently demonstrated that both ALDH and ADH domains in AdhE and BdhE possess shared enzymatic activities.

**Figure 2.**
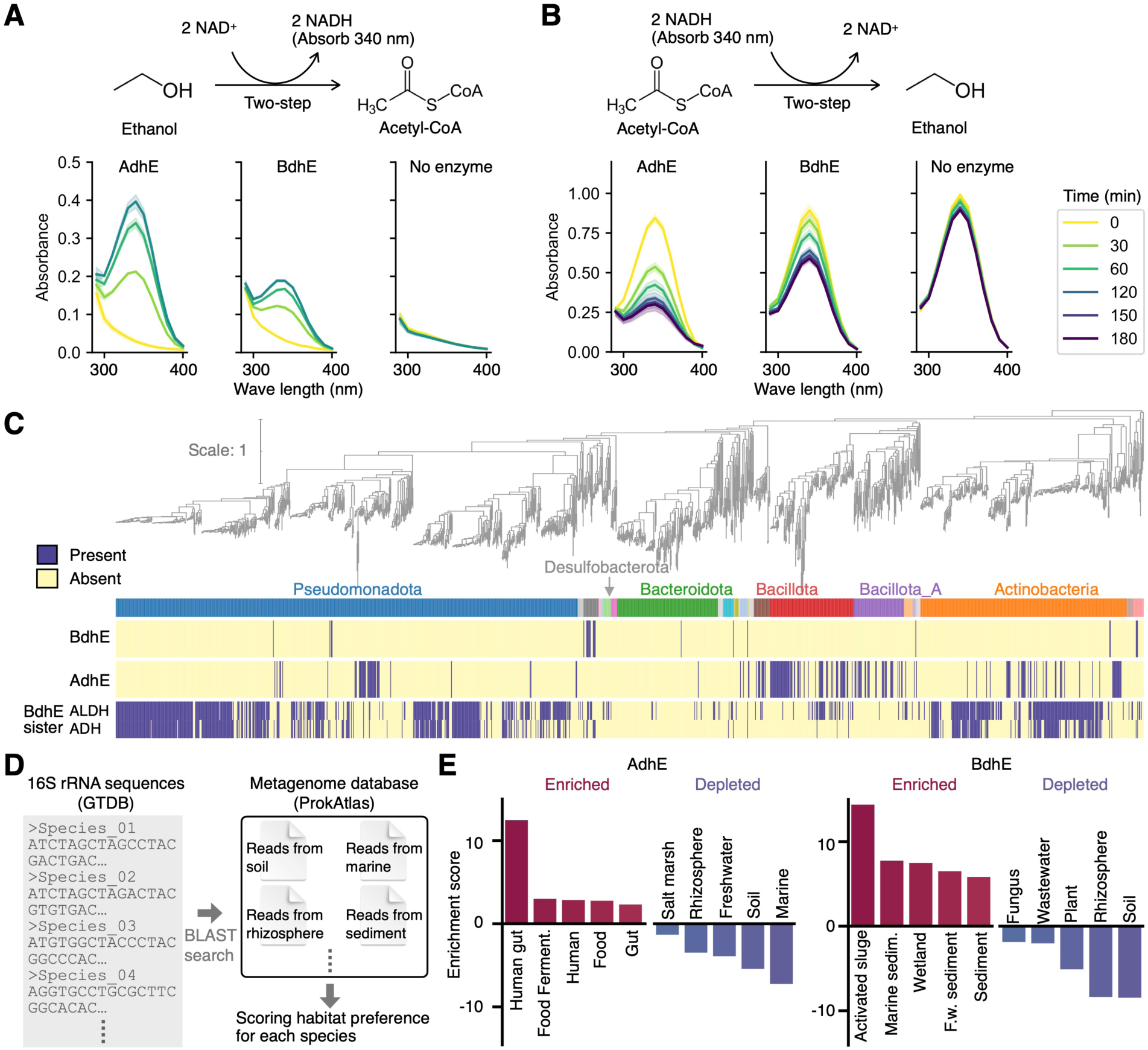
**Shared enzymatic activities and non-overlapped phylogenetic and ecological distributions of AdhE and BdhE**. **(A, B)** Assay results of ethanol oxidization activity (A) and Acetyl-CoA reduction (B) for AdhE and BdhE purified by a His-tag affinity column. The progression of the reactions was monitored by NADH-specific absorbance at approximately 340 nm. The yellow-to-purple curves were time-course absorbance spectra for conditions with AdhE, BdhE, or no enzyme. **(C)** Phylogenetic distribution of AdhE, BdhE and sister-clade gene families of BdhE. The species phylogenies contain all 43 species with BdhE, all 638 species with BdhE- sisters and 753 species out of 25,877 species with high-quality representative genomes. See **fig. S4** for the distribution on full species phylogeny. **(D)** The schematic diagram of habitat annotation for bacterial species based on 16S rRNA sequences of each species. **(E)** The habitat enrichment of species with BdhE or AdhE. The enrichment score represents the difference between each environment’s average habitat preference scores across species with and without AdhE/BdhE. The habitat preference scores were calculated using the ProkAtlas pipeline (*35*). Only top/bottom-five environments are shown. See also **fig. S5** for the full results. F.w. and sedim. stands for fresh water and sediment, respectively.

To explore the functional relationship of AdhE and BdhE in a macroevolutionary context, we compared their phylogenetic distributions across bacteria (**Fig. 2C**), as functionally coupled/redundant genes are known to co-occur/anti-cooccur in genomes in general, respectively (*32*, *33*). Both gene families exhibited broad yet sparse distribution across multiple phyla, suggesting the genes have been horizontally transferred across distant clades. Interestingly, none in the analyzed 25,877 species with high-quality complete genomes had both AdhE and BdhE. Furthermore, AdhE and each sister-clade ortholog of BdhE were found to show significantly complementary distribution by EvolCCM (*34*) (P < 2.22 ×10^-16^, **Fig. 2C, fig. S4A, B**), while the sister-clade families significantly tended to co-occur (P < 2.22 ×10^-16^), suggesting the two BdhE-sister families are functionally coupled and involved in redundant functions to those of AdhE. These results consistently suggest that the AdhE and BdhE are functionally redundant, and it is not adaptive to have both.

Given the non-overlapping phylogenetic distributions of *adhE* and *bdhE*, we hypothesized that these genes were distributed in distinct environments. We employed a pipeline to search 16S rRNA gene sequences in metagenomic datasets from diverse environments and scored habitat preferences for each species (*35*) (**Fig. 2D**). A comparison of habitat preferences revealed that *bdhE* is enriched in water-soil mixture environments such as activated sludge and marine/freshwater sediments, while *adhE* is enriched in human gut and other human-associated environments (**Fig. 2E and fig. S5**). These environments are commonly known to be anaerobic or to harbor anaerobic microbes (*36–38*), suggesting both AdhE and BdhE ecologically contribute to alcohol fermentation in anaerobic conditions, supporting our hypothesis of similar functions in distinct niches. Notably, BdhE was found in saline-associated environments (“marine sediment” and “marine”). Given choline oxidization activity was observed for BdhE, BdhE can also be involved in the synthesis of betaine, a well-known compatible solute for resistance in high-osmotic pressure environments (*39–41*). Overall, our results suggest that BdhE can act for similar or redundant functions to AdhE, while it is still possible that each family also have specific ecological roles in different environments.

### BdhE and AdhE evolved similar dimeric structural units forming distinct multimers

As both BdhE and AdhE showed shared enzymatic activities, we further investigated whether BdhE have evolved a similar 3D multimeric structure to that of AdhE following independent gene fusions. We first verified if both proteins form homocomplex structures by size-exclusion chromatography and blue-native polyacrylamide gel electrophoresis (BN-PAGE). Unexpectedly, BdhE showed a later retention time than AdhE, suggesting that the BdhE complex, if formed, has a smaller molecular size than the AdhE complex (**fig. S6A**). In addition, a BN-PAGE for BdhE resulted in a single band around the molecular weight of tetramers (391.6 kDa) (**fig. S6B**). Therefore, BdhE was suggested to form a complex, but it is not polymer like AdhE but a stable tetramer. Notably, the BN-PAGE for AdhE showed bands around the monomer and dimer weights, which is consistent with previous reports (*42*), probably because AdhE polymers were trapped at the starting point of the gel and only disassembled molecules were observed as bands.

As BdhE was suggested to form a homotetramer, we solved its quaternary structure using cryo-electron microscopy (cryo-EM), as well as that of AdhE (**Fig. 3A; fig S7**; gold-standard Fourier shell correlation (GSFSC) resolution of AdhE: 2.87 Å, BdhE: 2.55 Å). The results revealed that BdhE formed a ring-shaped tetramer, while AdhE formed a helical polymer, even though they were purified in the same condition. Two different conformations (extended and compact) have been reported for AdhE, and the structure we solved resembled the compact conformation, consistent with previous reports that AdhE forms compact conformation without cofactors (*26*, *28*). Negative stain images also supported the structural difference (**Fig. 3B and fig. S6C**). However, through modeling the structures of BdhE and AdhE, we revealed that both helical and ring-shaped structures consist of similar structural units of dimers (**Fig. 3C**). Both the BdhE and AdhE dimers showed a bent shape whose bending angles were different (**Fig. 3D and fig. S8A**). The slight structural difference likely contributes to the substantially distinct multimers, as the larger bending angle in AdhE compared to BdhE may prevent single-turn circle formation and instead promote the assembly of open-ended helical structures. It is also mentionable that AlphaFold2 (*43*) accurately predicted AdhE/BdhE homodimer units and even the tetramer structures of BdhE (**fig. S6D**). However, BdhE’s predicted circular structure might have been interpreted as an artifact without cryo-EM results, because AlphaFold2 preferred predicting ring structures even for AdhE, failing to predict the helical structures.

**Figure 3.**
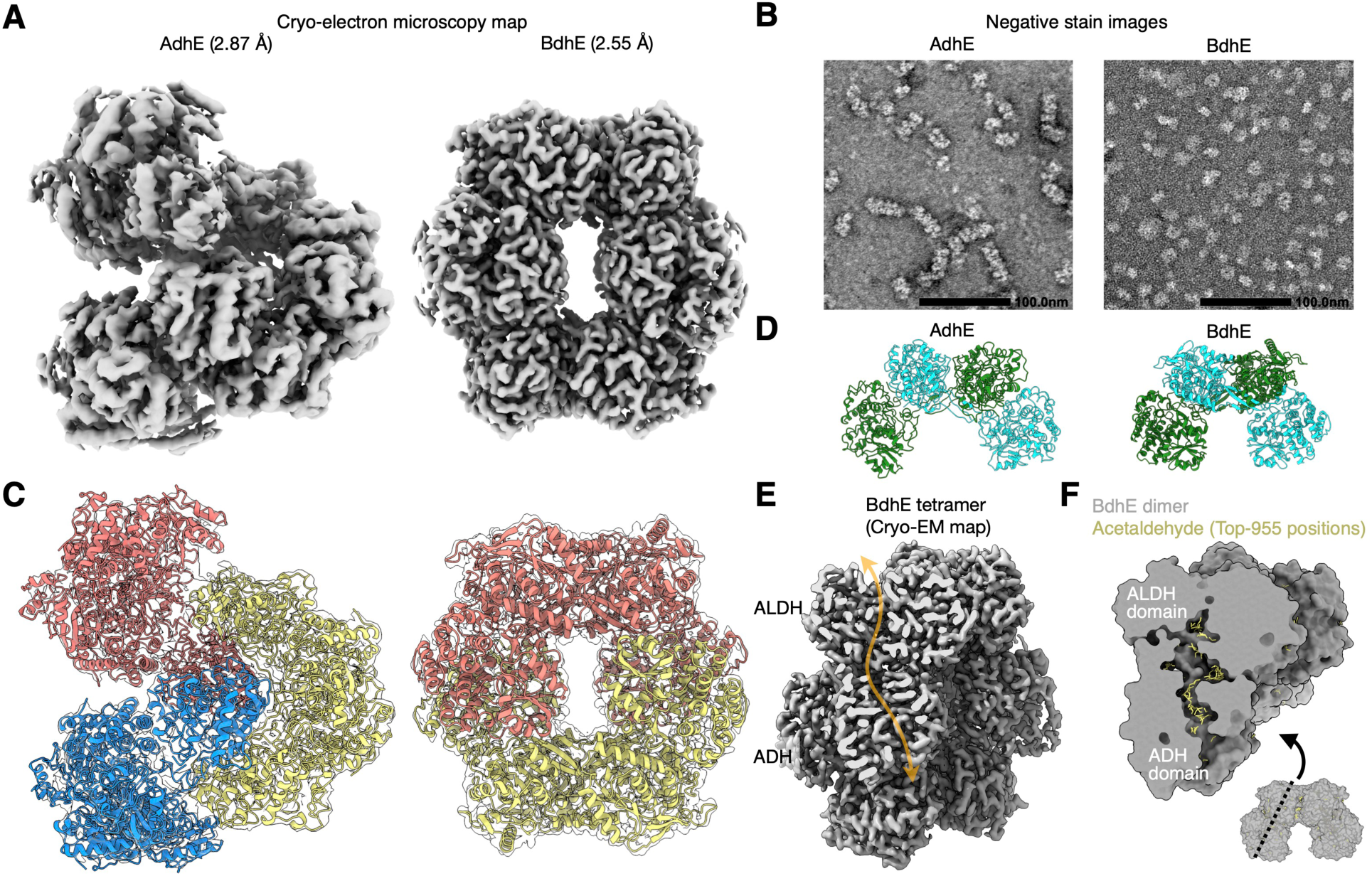
Cryo-electron microscopy structures of AdhE and BdhE suggesting convergence of inter-domain interactions. **(A)** The density maps of AdhE and BdhE. **(B)** The negative stain electron microscopy analysis of AdhE and BdhE. Scale bars represent 100.0 nm. The image is trimmed to fit in a space. See also **fig. S6C** for the full images. **(C)** The structural models of AdhE and BdhE superimposed with electron density maps. The red, yellow, and blue parts represent different dimeric structural units. **(D)** The structure of dimeric structural units of AdhE polymers and BdhE tetramers shown in (C). Green and cyan molecules represent different protein molecules. **(E)** A cross-section of BdhE tetramer where ALDH and ADH domains were closely positioned. The transparent orange path represents the tunnel through which substrates pass. **(F)** Docking simulation results of a BdhE dimer and an acetaldehyde molecule. The yellow molecular structures represent 955 possible docking positions of acetaldehyde within 4 kcal/mol from the lowest-energy position.

### BdhE and AdhE convergently evolved channeling interfaces via distinct loop elongation

In the AdhE and BdhE dimers, the proximity between ALDH and ADH domains from different protein molecules was commonly observed, suggesting that BdhE exhibits substrate channeling similar to AdhE by forming the dimeric structure. Looking into the cross-section of the BdhE tetramer’s electron density, we found that the tunnels where substrate passes were connected between ALDH and ADH domains (**Fig. 3E**). Similarly to a previous study of AdhE (*28*), we further confirmed that a substrate molecule could be inside the tunnel stably by docking simulation both in AdhE and BdhE dimers (**Fig. 3F and Fig. S8B**), supporting the possibility that both fusion proteins have convergently evolved substrate channeling by independently acquiring ALDH-ADH contacts.

To characterize how AdhE and BdhE evolved the ALDH-ADH contact interfaces, we compared the interaction interface residues in their dimeric structures (**Fig. 4A**). We found the ALDH-ADH interface was involved with multiple loop structures (loop 1 and 2 of AdhE and loop 3 and 4 of BdhE; **Fig. 4B**), which is consistent with a previous report regarding AdhE (*26*). Intriguingly, these interface-forming loops of AdhE and BdhE were non-homologous, and the inter-domain interactions were achieved by AdhE- and BdhE-specific manners, respectively. Furthermore, we found that these loop sequences were AdhE- or BdhE-specific and absent even in the sister-clade ALDH or ADH genes of AdhE and BdhE (**Fig. 4C**), suggesting AdhE and BdhE repeatedly evolved channeling interfaces using independently elongated distinct loop structures. Therefore, AdhE and BdhE convergently evolved inter- domain interactions at structure level, rather than at residue level.

**Figure 4.**
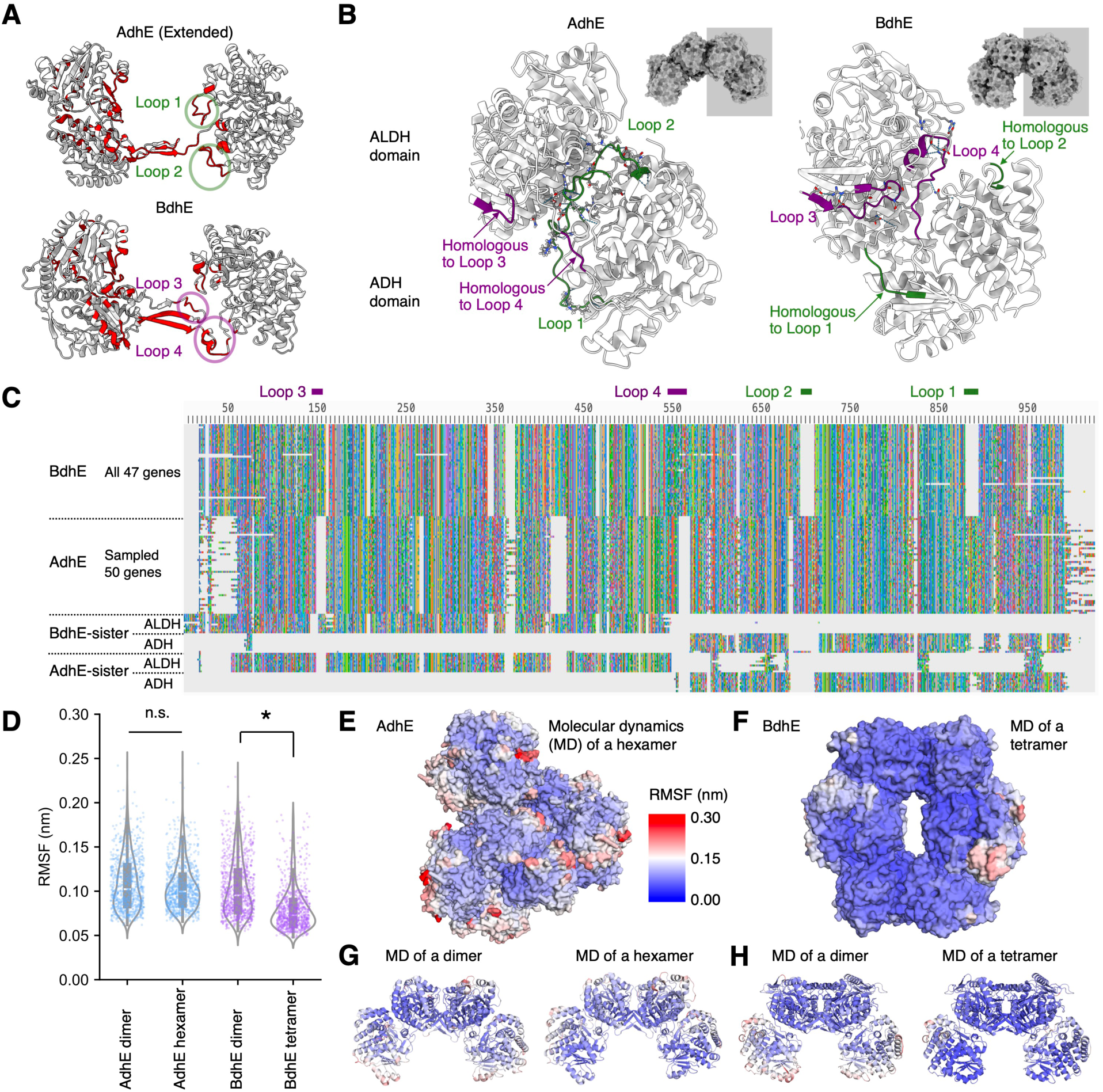
Distinct properties of convergently evolved AdhE and BdhE multimers. (A) **The** residues involved in inter-molecule interactions of AdhE and BdhE dimers. The red residues represent the interaction interfaces estimated by PICKLUSTER (without-clustering mode) implemented in ChimeraX-v1.7rc202311290355. The AdhE dimer structure of the extended conformation was retrieved from the Protein Data Bank (PDB; ID: 6TQH). Loops 1-4 were annotated to the particular loop structures where inter-molecule interaction residues of AdhE or BdhE enriched. **(B)** The ALDH-ADH interaction interface of AdhE and BdhE. The green and purple loop structures represent the loop 1-4 and their homologous parts. **(C)** Multiple sequence alignment of BdhE, AdhE, and sister-clade single-domain families of them. All 47 BdhEs, randomly sampled 50 AdhEs, and randomly sampled 10 genes for each of the other families are shown. The positions corresponding to loops 1-4 are indicated by green and purple bars. **(D)** Comparison of RMSF distributions among dimers and hexamer/tetramer of AdhE and BdhE, respectively. RMSFs were calculated for each alpha-C atom of every residue. The RMSFs of an AdhE hexamer were calculated for the intermediate dimer unit. The asterisk indicates the statistical difference (Two-sided Wilcoxon signed rank test. *p =* 0.11 and 0.00 for AdhE and BdhE, respectively.). **(E-H)** Color mapping of RMSF onto the 3D structures of an AdhE hexamer (E), a BdhE tetramer (F), an AdhE dimer (G), and a BdhE dimer (H). In Fig. 4G and H, RMSFs calculated by molecular dynamics (MD) for dimer structures are shown on the left, and RMSFs for hexamer/tetramer structures mapped onto dimer structures are shown on the right, respectively.

The difference in channeling interfaces of AdhE and BdhE resulted in structural differences in the tunnel where the substrates would be passed. We found a gap in the tunnel wall formed by a BdhE dimer, which were not found in AdhE dimers (**fig. S8C-E**). In an AdhE dimer, the linker of ALDH and ADH domains filled the gap, while the linker of BdhE formed the elongated loop (loop 2 in **Fig. 4A and B**) and thereby remained the gap. In BdhE, however, another BdhE dimer partially filled the gap in the tunnel wall within the tetrameric complex (**Fig. S8D**), suggesting that the tetrameric structure is more effective at preventing intermediate aldehydes from leaking out than the dimeric structure.

### The difference between helical and circular multimer structures is associated with structural stability

As the similar dimeric structural units of AdhE and BdhE form distinct multimeric structures (i.e., helical polymer of AdhE and circular tetramer of BdhE), we further hypothesized the overall structural difference leads to different features in terms of structural stability. To test this, we conducted molecular dynamics (MD) simulation of an AdhE hexamer, a BdhE tetramer, and their dimers for 5 ns, which is long enough to see the structural fluctuations at stable states (**fig. S8F**). To assess the structural stability, root mean square fluctuations (RMSFs) were calculated for each residue over the period from 1 ns to 5 ns (**Fig. 4D** and **fig. S8G**). For the AdhE hexamer, we calculated RMSF for the dimer not locating at helix terminals. The results showed that the BdhE tetramer showed decreased RMSF, or increased the structural stability, compared to BdhE dimer, while AdhE hexamer formation did not increase the stability compared to AdhE dimer (**Fig. 4D**). By mapping the RMSF values onto the 3D structures of AdhE and BdhE, we found the BdhE showed higher structural stability than the AdhE across the whole structure (**Fig. 4E and F**). We further found that the stabilization by forming tetramers of BdhE was observed for most parts of the structures (**Fig. 4G and H**). These results indicate that the tetramer formation of BdhE contributes to higher stability than forming dimers, while polymer formation of AdhE does not, possibly due to the flexibility difference between circular and helical structures.

### Genomic neighboring of ALDH and ADH genes underlies repeated gene fusions

As distinct pairs of ALDH and ADH genes repeatedly fused into *adhE* and *bdhE*, we next asked whether any common genomic background facilitated fusions of those genes. Gene fusions often occur between genes neighboring in genomes (*15*), so we hypothesized that single-domain ALDH and ADH genes tend to be neighbors in bacterial genomes. We comprehensively detected neighboring gene pairs located within ten genes for various ALDH and ADH orthologs across 26,778 high-quality representative genomes of prokaryotes and found that neighboring genes are enriched in specific ALDH and ADH ortholog pairs (**Fig. 5A**). The adjacent gene pairs include *pduP*-*pduQ* (KEGG ID: K13922-K13921), *eutE*-*eutG* (K04021-K04022), and *gbsA*-*gbsB* (K00130-K11440), which have been reported to be coded in the same operons and to be functionally coupled (*40*, *44*, *45*). These results suggest that functionally coupled ALDH and ADH genes repeatedly become neighbors in genomes and are transcribed together in the same mRNA to adjust the expression levels of enzymes for a series of metabolic reactions.

**Figure 5.**
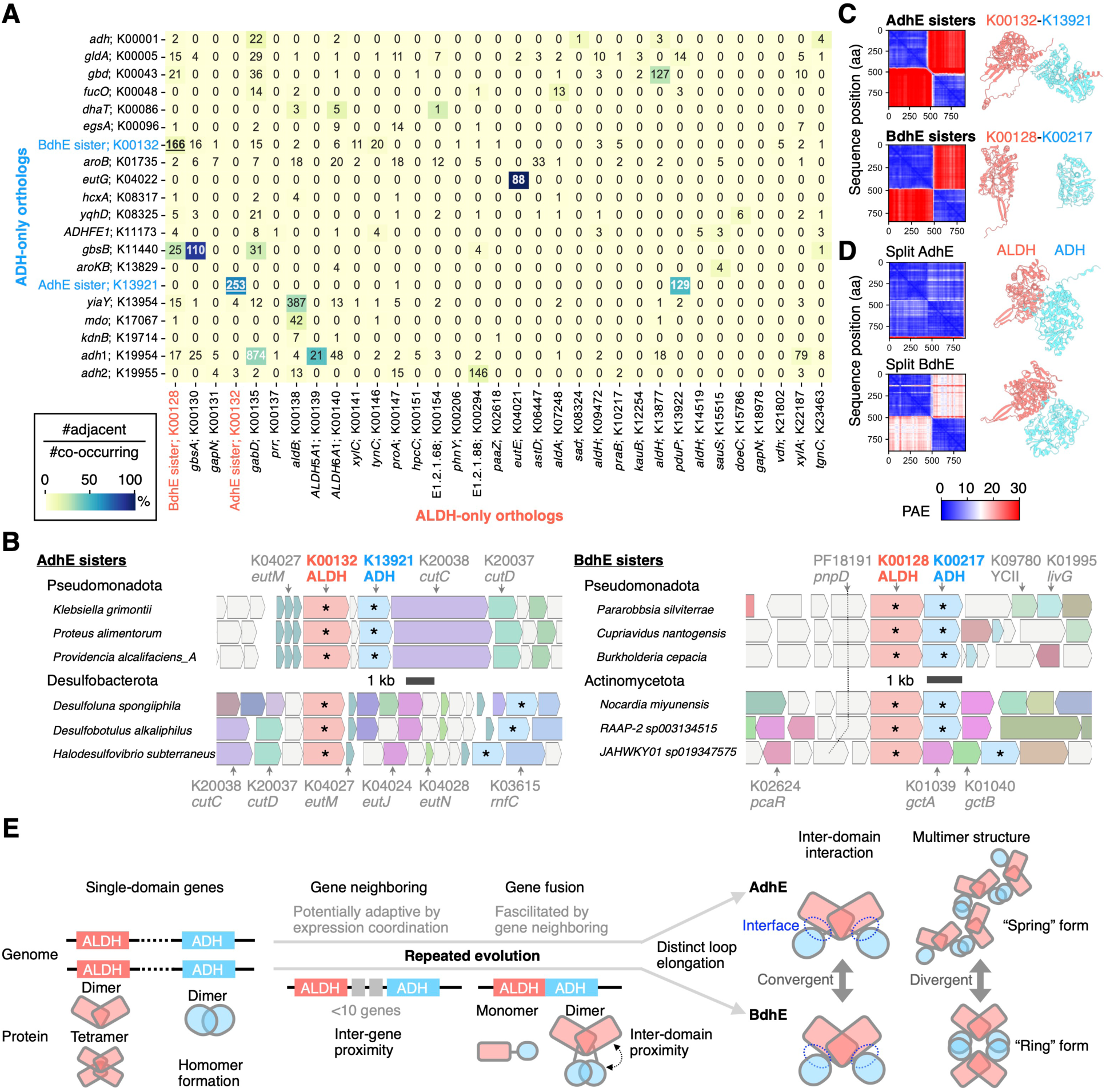
Genomic evolution preceding recurrent gene fusion events and the model for repeated evolution of AdhE and BdhE. **(A)** A heatmap showing the counts/ratios of the pairs of ALDH and ADH genes adjacently coded in prokaryotic genomes, i.e., coded in the same strand of the same contigs intervened by <10 genes. The annotated numbers indicate the gene pair counts in 26,778 prokaryotic genomes, and the color scale represents the ratio of the number of adjacent gene pairs per total number of pairs co-occurring in the same genomes. The ALDH and ADH orthologs in this panel correspond to ALDH/ADH orthologs included in the phylogeny of Fig. 1C **and D**. The number of gene pairs of AdhE- and BdhE-sister orthologs are underlined. **(B)** Multi-species comparison of gene synteny around AdhE- and BdhE-sister orthologs. Here we sampled three species in different genera for two phyla, whose genomes have AdhE- and BdhE-sister orthologs as neighbors (genes with asterisks). The colored genes are assigned KEGG ortholog identifiers. The visualization was conducted by Annoview version 2.0 (*47*). **(C)** ColabFold prediction of heteromeric interaction between AdhE-sister or BdhE-sister ALDH and ADH proteins adjacently coded in genomes. Structural predictions were conducted by ColabFold v1.5.2. Here, we show the results of an example protein pair for the AdhE and BdhE sisters, respectively. The left heatmap and the right structures indicate the positioned aligned errors (PAEs) and the predicted structures where ALDH and ADH proteins are colored in red and cyan, respectively. **(D)** ColabFold structure prediction of heteromeric interaction between split AdhE/BdhE domains. Here, we show the results for AdhE and BdhE used in our enzymatic and structural analysis. **(E)** A schematic model of repeated genomic and structural evolution leading to extant AdhE and BdhE.

Based on phylogenies of ALDH and ADH domains (**Fig. 1C and D, Fig. S2D**), we have identified pairs of orthologs, K00132-K13921 and K00128-K00217 as ALDH- and ADH-only sister clades of AdhE and BdhE, respectively. By analyzing the neighboring gene pairs, we realized that the sister-clade families of AdhE/BdhE tend to be coded by neighbor genes, respectively (**Fig. 5A**). The synteny around these genes were conserved and they were adjacent to genes with potential functional relationships with ALDH/ADH genes (**Fig. 5B**). For example, the pairs of AdhE sisters were close to *eutM* coding major elements of bacterial microcompartments where ethanol metabolism is catalysed (*46*), and BdhE sisters were also close to *gctA/B* (gluconate CoA transferase) which may contribute to produce carbonyl-CoA as substrates of an ALDH. The result suggests the ancestral single-domain genes of both *adhE* and *bdhE* were originally coded in genomically close positions before gene fusion and thereby enabled to fuse by small deletions. Although the functions and expressions of the sister-clade families were seemingly coupled, the adjacently coded proteins were not predicted to show inter-domain interactions, and the results were robust for any other pair of ALDH and ADH families (**Fig. 5C, fig. S9A**). In contrast, split domains of AdhE and BdhE as positive controls were predicted to interact with each other (**Fig. 5D**). Therefore, the convergent ALDH-ADH interactions observed in AdhE and BdhE were suggested to have evolved after gene fusions through acquisitions of the family-specific loops (**Fig. 4A**). Notably, all those single-domain proteins were predicted to form homodimers whose interaction interfaces resembled ALDH-ALDH and ADH-ADH interfaces in both BdhE and AdhE (**fig. S9B, C, and S10**), suggesting the helical or circular multimers of AdhE and BdhE were achieved by both homomeric interactions inherited from ancestors and convergently acquired ALDH-ADH interactions.

## Discussion

Consequently, we identified BdhE, an ALDH-ADH protein family that emerged by gene fusion independently of AdhE. We further showed that AdhE and BdhE convergently evolved ALDH-ADH interactions with distinct loop elongations while their multimer structures highly diverged (a helical polymer and a circular tetramer). Based on these findings, we propose a model of AdhE/BdhE evolution (**Fig. 5E**), in which the repeated evolution of AdhE and BdhE would have started from distantly positioned genes coding ALDH and ADH enzymes which form homodimers and/or homotetramers (*48*) (**fig. S9B, C**). Then, some pairs of the genes became neighbors in genomes (**Fig. 5A**), and the neighbor genes were fused (**Fig. 1B-D**). The fused protein would show only ALDH-ALDH and ADH-ADH interactions at first (**Fig. 5B**). Then, the fused enzyme families convergently evolved elongated loops and interactions between ALDH and ADH domains, suggested to have enabled substrate channeling (**Fig. 3E and F**). As a result of the inter-domain interaction, AdhE evolved spring-like helical polymers with variable pitch lengths while BdhE evolved ring-like tetramers, both using the interaction interfaces for newly evolved ALDH-ADH contacts and ancestral homomer formation (**Fig. 3A-C**). In this way, repeated gene fusions led to convergence of inter-domain contacts and dimeric structures while divergence of overall multimer structures.

This stepwise repeated evolution of AdhE and BdhE suggests evolutionary patterns of protein structures and underlying their principles. Unlike well-documented functional convergence between distinct protein families and residue-level convergence within families (*6*, *8*), our study provides a model case of structural convergence through the repeated evolution of inter-domain and inter-molecular interactions. Our findings suggest three evolutionary patterns. First, gene neighboring precedes fusion protein evolution. This pattern can be common across many different fusion gene families (*15*), as the neighboring genes can be co-regulated and horizontally transferred together within operons. In the case of ALDH and ADH, the coordination of gene expression levels can be adaptive by preventing the accumulation of cytotoxic aldehyde as intermediates. Proximities of genes in genomes can also facilitate gene fusions by reducing the fitness loss of deletion achieving the fusions.

As the second pattern, gene fusions precede structural convergence of inter-domain interactions.

Gene fusion can fix the proximity of previously uncontacted domains and stabilize incomplete inter- domain interactions to facilitate the following inter-domain contact evolution. While a previous study proposed domain interaction evolution precedes gene fusions (*49*), the case of AdhE and BdhE suggests the opposite order is also possible especially when the gene fusions are facilitated by gene neighboring. Note that AdhE and BdhE showed similar orientations of interacting ALDH and ADH domains, while substrate channeling could be achieved by other orientations (i.e., rotating a domain keeping the substrate channel), which may suggest the particular orientation of domains were favored during evolution because of any evolutionary constraints such as thermostability of the protein structures (*50*).

As the third pattern, structural innovation during evolution such as inter-domain interactions are achieved by a newly elongated loop, as reported in the evolution of enzymatic activities, too (*51*).

Existing structures constrain evolutionary changes, so that evolution tends to favor alterations in loop structures that don’t disrupt existing structural elements. These three repeated and explainable evolutionary patterns provide us clues to predict the evolution of existing proteins. For example, we can predict which ALDH and ADH genes will likely fuse in the future based on the current gene neighboring states (**Fig. 5A**), and what structural evolution follows the gene fusions.

Our study also shed light on bacterial adaptation strategies to mitigate aldehyde toxicity, which is universal for diverse organisms including humans (*52*, *53*). The results suggest neighboring of ALDH and ADH genes in genomes and ALDH-ADH channeling to prevent aldehyde leakage from enzymes as versatile aldehyde mitigation mechanisms in bacteria. Some of the neighbor ALDH and ADH genes code enzymes known to be confined in metabolosomes, which are also thought to act as aldehyde mitigation (*46*, *54*), suggesting multiple distinct mechanisms have been evolved to avoid aldehyde accumulation in bacterial metabolic evolution. Substrate channeling within molecules of AdhE or BdhE can be a more cost-effective strategy than producing the massive apparatus of metabolosomes.

While the ALDH-ADH interaction repeatedly evolved, AdhE and BdhE showed distinct quaternary structures (i.e., “spring” and “ring” form). The finding suggests that the whole multimeric structures can substantially diverge by slight difference of structural units, such as the bending angle of dimeric units. The spring-ring differences between the two families potentially result in the functional properties of the enzymes. While spring-form AdhE has been suggested to regulate its activity depending on cofactor binding by dynamically changing the helix pitch lengths (*26*, *28*), BdhE forms a ring structure which will not show such structural freedom observed in helical structures (i.e. dynamic pitch length) (**Fig. 4D-H**). The ring structure of BdhE is thus suggested to confer more rigid enzymatic activity than AdhE against environmental changes (e.g., salinity, pH, or temperature). Therefore, the spring-ring difference can be caused by distinct selective pressures for flexibility or stability of enzymatic functions, although it could also be a result of neutral evolution. Indeed, species with BdhE inhabit diverse natural environments, including marine, freshwater, hydrothermal vents, and hot springs, although those with AdhE mainly inhabit human-associated stable environments (**Fig. 2E and fig. S6**). As proteins showing rigid structures are generally found in thermophiles (*55*), the multimeric structural difference between AdhE and BdhE can be associated with ecological niche difference.

Several protein families, including Rad51 (*56*), MuB (*57*), CK2 (*58*), and allophycocyanin (*59*), exhibit both helical (or linear) polymer and circular multimer structures, with functional differences reported between these conformations. The helical-circular transitions in allophycocyanin, analogous to the AdhE-BdhE structural divergence, stem from variable bending angles of structural units, suggesting a general molecular mechanism leading to multimeric structure diversity across protein families. Rad51, MuB, and CK2 were shown to have conformational plasticity within individual molecules, and allophycocyanin transitioned between stable conformations during evolution within the protein family (*56–59*). The evolution of AdhE and BdhE, on the other hand, represents a unique case of helical-circular divergence by convergent evolution of inter-domain interactions in independently emerged fusion proteins. This convergent evolution revealed in this study offers valuable insights for exploring alternative amino acid sequence spaces that encode similar inter-domain interactions, hinting the design of proteins with comparable monomer structures but orthogonal multimeric assembly into ring and spring conformations. It is also noteworthy that AdhE and BdhE may have undergone transitions between different assembly states (helical and circular conformations) within each family, considering recent evidences showing protein families experience changes in their quaternary structure during evolution (*59*, *60*).

In this study, we demonstrated a clear example of macroscopic protein structure convergence, where similar dimeric structures evolved through independently acquired interactions. This finding extends our understanding of protein convergence beyond residue and domain levels to inter-domain and inter-molecular levels. Further investigation of protein structural evolution, particularly in proteins with convergent domain architectures, may uncover hidden evolutionary repeatability of inter-domain interactions. The latest rapid expansion of protein structure universe—driven by advances in cryo- electron microscopy technologies and structure prediction tools like AlphaFold and ESMFold (*43*, *61*, *62*)—has opened unprecedented era for elucidating the patterns and principles underlying protein structural evolution.

## Methods

### Dataset

We retrieved all the protein sequences of every prokaryotic representative genome and reference phylogenies from GTDB r202 on April 28, 2021. The datasets contained 45,555 bacterial and 2,339 archaeal species of all the phyla defined in GTDB. We annotated orthologs for each gene in all the representative genome based on KEGG Orthology by KofamScan v1.3.0 (*1*). We also extracted a tree of all the 25,877 bacterial species for which high-quality representative genomes (defined as >95% completeness and <5% contamination throughout this study) are available in GTDB, using the TreeNode.prune function (“preserve_branch_length = True”) in ete3 toolkit 3.1.2 (*51*). 16S rRNA gene sequences for 22,304 of the 25,877 species were also downloaded from GTDB r202.

### Extraction of non-AdhE ALDH-ADH fusion proteins

As described previously (*2*), we conducted sensitive sequence similarity search of AdhE against all the proteins coded in 45,555 bacterial genomes from GTDB using MMseqs v13.45111 (easy-search -s 7.50), by querying 4,833 proteins annotated as AdhE (K04072) by KofamScan and showing >800 aa length in GTDB genomes. Because the sequence similarity search suggested existence of non-AdhE ALDH-ADH fusion proteins, we next retrieved all the 8,472 protein sequences with ALDH (PF00171) and ADH domain (PF00465) from InterPro (retrieval date: November 29, 2022), in which 8,467 and 5 proteins had ALDH-ADH and ADH-ALDH domain architecture, respectively. After extracting ALDH-ADH proteins which include AdhE, we annotated KEGG Orthology identifiers by KofamScan v1.3.0. Then, we extracted 23 proteins not annotated as AdhE (K04072 in KEGG Orthology) and with >800 aa length as non-AdhE ALDH-ADH proteins. We next searched their homologs in bacteria by querying N-terminal 480 aa (corresponding to ALDH domains) or C-terminal 370 aa (corresponding to ADH domains) of those 23 proteins against all the proteins coded in 45,555 bacterial genomes from GTDB. The sequence searches resulted in 7,370,016 and 1,718,637 proteins hit by querying the N-terminal and C-terminal sequences, respectively. From the hits, we extracted non-AdhE ALDH-ADH proteins which showed >40% sequence identity for both N-terminal and C-terminal queries. We also obtained single-domain close homologs of the non-AdhE ALDH-ADH proteins by extracting hits showing >50% and >40% alignment identity for N-terminal and C-terminal queries, respectively. Lastly, we aligned those ALDH-ADH proteins and their close homologs by MAFFT v7.310 (no option), and added an AdhE sequence (from the *E. coli* genome in GTDB) to the alignment as an outgroup by MAFFT (--add). We next trimmed alignment columns dominated by gaps with TrimAl v1.4rev15 (-gappyout). Then, we constructed gene phylogenies for both protein domains by IQTree v2.0.3 (-m MFP -bb 1000 -nt 20). We extracted 47 proteins showing >700 aa as BdhE and confirmed their monomer structures predicted by AlphaFold2 are all similar to each other excluding two proteins with unidentified amino acids. We named the extracted ALDH-ADH fusion proteins as BdhE (Bifunctional dehydrogenase E), a new fusion enzyme family.

### Phylogenetic analysis of AdhE and BdhE across diverse ALDH and ADH proteins

To unveil the phylogenetic relationship between AdhE and BdhE among diverse ALDH and ADH protein families, we firstly extracted KEGG Orthologs annotated for genes coding ALDH (Pfam ID: PF00171) or ADH domains (Pfam ID: PF00465 (“Fe-ADH”) or PF13685 (“Fe-ADH_2”)) in KEGG Genes database (retrieval date: March 31, 2022). We excluded 13 of the extracted orthologs (K00254, K00891, K01999, K01588, K01647, K03106, K03110, K03469, K03601, K07029, K13821, K13829, and K13830 in KEGG Orthology), because they coded non-enzymatic genes and/or false positives caused by genes fused with other ALDH or ADH genes. The ALDH- and ADH-only KEGG orthologs after the curation are shown in **Fig. 5A**. Next, we collected protein sequences of randomly selected 10 genes in GTDB bacterial genomes for each of the KEGG Orthologs. Then, we merged the protein sequences of single-domain orthologs, randomly sampled 200 AdhEs, and all the 47 BdhEs, and reconstruct the domain-wise gene phylogenies. Multiple sequence alignment, trimming, and phylogeny estimation were done by MAFFT v7.310 (-- thread 20), Trimal v1.4rev15 (-gt 0.7), and IQTree v2.0.3 (-m MFP -bb 1000 -nt 10). To test if AdhE and each of BdhE sister families (K00128 and K00217) significantly tend to be co-occurring or anti- cooccurring in genomes, we used EvolCCM (*3*) to construct a model for gains and losses of AdhE, K00128, and K00217 across the reference genome phylogeny.

### Habitat preference analysis of whole bacterial species

To investigate environments where species with AdhE or BdhE enriched in, we estimated the habitat preference for each of the 22,304 bacterial species for which high-quality representative genomes and 16S rRNA sequences were available from GTDB (see Datasets). We queried the 16S rRNA sequences of the 22,304 species against ProkAtlas online (*4*) on June 30, 2021. ProkAtlas evaluates habitat preferences by comparing 16S rRNA sequences to short-read metagenomic data from various environments. Based on these results, ProkAtlas assigns a habitat preference score for each environment for every species. We used BLAST searches with specific criteria: 97% nucleotide identity and a minimum of 150 bp coverage.

### Expression and Purification of AdhE and BdhE

The full-length *adhE* gene from *Escherichia coli* (*E. coli*) K12 strain (UniProt ID: P0A9Q7) and the full-length *bdhE* gene from the *Halomonas eurihalina* genome in GTDB were artificially synthesized with codons optimized for expression in *E. coli.* Each gene was fused with an N-terminal 6-His tag followed by a thrombin cleavage site (LVPR|GS) and inserted into the pET28a (+) vector (Twist BioScience)).

Both AdhE and BdhE were expressed in BL21 (DE3) cells by induction with 1 mM isopropyl β-D-thiogalactopyranoside (IPTG) at 18 °C overnight after the preculture until the OD reached 0.4-0.5 at 37 °C in LB medium. Then, we harvested cells by centrifugation at 5,000 rpm 4 °C for 15 min, resuspended and sonicated in a lysis buffer (buffer A) containing 50 mM Tris-HCl pH 8.0, 500 mM NaCl, and 5% (v/v) glycerol with 2 mg/mL lysozyme. We clarified the sonicated lysate by centrifugation at 15,000 g, 10 min and 4 °C and filtration with 0.2 μm filter. Then, we conducted His-tag purification using TALON Spin Columns purification (Takara, Cat. No. 635603). We washed the TALON beads with HisTALON Equilibration buffer or Buffer A, and loaded the filtered lysate after sonication. Then, we washed the beads with buffer A containing 50 mM imidazole. The bound AdhE or BdhE proteins were eluted by buffer A containing 100, 150, and 200 mM imidazole. We removed imidazole from the eluates by dialysis in 100 mM NaCl, 1 mM DTT, and 0.5 mM EDTA at 4°C overnight. We further purified AdhE and BdhE by gel-filtration chromatography column (Cytiva; 28-9909-44; Superdex 200 Increase 10/300 GL) equilibrated with buffer containing 50 mM Tris-HCl pH 8.0 and 100 mM NaCl. AdhE- or BdhE- containing fractions were collected and further concentrated with Amicon Ultra30K MWCO centrifugal filters (Millipore; UFC903024) up to >4 mg/mL. The purified samples were flash-frozen for storage in liquid nitrogen before storage at -80 °C. For proteins used for cryo-electron microscopy (cryo-EM) experiment, we did not freeze the purified protein samples until the cryo-EM sample preparation.

### Enzymatic activity assay for AdhE and BdhE

To investigate the enzymatic activities of AdhE and BdhE, we quantitated the decrease or increase of absorbance by nicotinamide adenine dinucleotide (NADH) peaked at a wavelength of ∼340 nm, referring to previous protocols for AdhE enzymatic assay (*5*). We used a plate reader (Tecan; Infinite 200 Pro) to measure absorbance spectra across 230-400 nm with two replicates (**Fig. 2A, B**), and to measure absorbance values at 360 ± 35 nm with four replicates (**fig. S4B, C**). All assays were conducted at 37 °C, and the total volume was 100 μL (**Fig. 2A, B**) or 40 μL (**fig. S4B, C**). In **Fig. 2A, B**, we measured the activities of the AdhE and BdhE for the forward reaction (oxidization of NADH) in a mixture containing 50 mM Tris-HCl pH 8.0, 50 μM FeSO4, 200 μM acetyl-CoA, and 500 μM NADH with 6 μg of AdhE or BdhE purified by the histidine tag. For the reversed reaction (reduction of NAD^+^), we assayed the activity in a mixture containing 50 mM Tris-HCl pH 8.0, 50 μM FeSO4, 200 μM CoA-SH, 500 μM NAD^+^, and 200 mM ethanol with 22 μg of AdhE or BdhE. In **fig. S4B** and **C**, while most conditions were the same as Fig. 2A and **B**, the concentration of substrate (acetaldehyde or acetyl-CoA) was 10 mM for forward reaction, and the amount of AdhE or BdhE was 1.8 μg (forward reaction) or 2.0 μg (reverse reaction), which was purified by the histidine tag and the gel-filtration chromatography.

### Sodium dodecyl sulfate- and blue native- polyacrylamide gel electrophoresis (SDS-PAGE and BN-PAGE)

We conducted SDS-PAGE using a 12.5% polyacrylamide gel at 21.0 mA for ∼45 minutes (ATTO; WSE- 1025). Products and intermediates of His-tag purification of AdhE and BdhE were loaded after adding loading buffer (ATTO; AE-1430) and heating at 100 °C for 5 minutes. The molecular mass of each band was estimated based on molecular weight markers (Broad; 3452). We also conducted BN-PAGE using 3- 14% polyacrylamide gradient (ATTO; UH-T314) at 150 V (constant V) for ∼75 minutes in a running buffer (ATTO; WSE-7057). Purified AdhE 5 or 10 μg and BdhE 1 or 2 μg were applied with loading buffer (ATTO; WSE-7011). The molecular mass of the AdhE/BdhE multimers was estimated based on molecular weight markers (ATTO; WSE-7016). A BN gel lane was destained in a water solution containing 50% methanol and 12.5% acetic acid, and then re-stained with EzStain Aqua (ATTO; AE- 1340).

### Negative stain experiments

To prepare grids for negative stain observation, we applied 3 μL of 50 ng/μL purified AdhE or BdhE to an elastic carbon-coated Cu (250 mesh) grid (Okenshoji; ELS-C10), and dried the grid with filter paper. Then, we stained the grid with 3 μL of 1% uranium acetate three times and dried the grid at room temperature. Excess uranyl acetate was eliminated by filter paper. Then, the prepared grids were observed using a JEM-1400Flash electron microscopy with sCMOS camera (Matataki Flash).

### Cryo-electron microscopy experiments, single particle analyses, and model constructions

To prepare grids for cryo-electron microscopy, we applied 3 μL of 2.02 mg/mL AdhE or 1.35 mg/mL BdhE to a Quantifoil holey carbon grid (R1.2/1.3, 300 mesh copper, Quantifoil Micro Tools GmbH),, which was hydrophilized by PIB-10 plasma ion bombarder (Vacuum Device) in advance. Using Vitrobot Mark Ⅳ (ThermoFisher Scientific), we blotted the protein samples with a setting of 10 blot force at 6 °C in 100% humidity and froze them immediately in liquid ethane, setting “wait time” and “blot time” as 3 and 4 seconds respectively. Then we collected micrograph images in movie mode using Talos Arctica G2 (ThermoFisher Scientific) with a K2 direct detector (Gatan) operated at 200 kV, 1.03 Å/pixel, 50 e^-^ /Å^2^/micrograph with a -1.75 to -1.0 μm defocus range and 50 movie frames. As shown in **Fig. S7C**, we conducted single particle analysis for AdhE and BdhE by CryoSPARC v4.4.1. Note that CTF refinement was performed simultaneously with all the homogeneous refinements in CryoSPARC. Molecular structure models were built with Coot v0.9.8.91 (*6*) started from a previously reported model (PDB ID: 6TQM) and an AlphaFold-predicted model of AdhE and BdhE, respectively. We then refined with the real-space refinement procedure implemented in Phenix v1.20.1-4487 (*7*). Electron density maps and molecular structure models were visualized by UCSF ChimeraX v1.7rc202311290355.

### Docking simulation

To conduct docking simulation and molecular dynamics, we prepared dimeric structure models of the extended and compact form of AdhE from the Protein Data Bank (PDB; ID: 6TQH and 6TQM) and the dimeric BdhE model determined by this study. The models were firstly aligned by Matchmaker algorithm implemented in ChimeraX v1.7rc202311290355. The proteins and acetaldehyde structural models for docking simulation were prepared using Openbabel v3.1.0 (*8*). and UCSF Chimera v1.17.3. The docking simulation was performed using AutoDock Vina v1.2.5 (*9*, *10*) with a box size of X = 140.0, Y = 100.0, and Z = 100.0. The parameters of exhaustiveness, number of modes, and energy range were set to 100, 1000, and 4, respectively.

### Molecular dynamics simulations

All molecular dynamics simulations were performed using the GROMACS (version 2023.2) molecular dynamics simulation package (*11*). The initial atomic coordinates of AdhE hexamer (compact form) was obtained from the PDB (ID: 6AHC), and the model of BdhE was constructed in this study. These structures were converted into topology and coordinate files utilizing GROMACS. The CHARMM36m force field was used for the protein (*12*), and the CHARMM General Force Field was used for the ligand (*13*). Water molecule was simulated with the TIP3P model (*14*). Sodium ions were added to neutralize each system. Energy minimization was performed using the Steepest Descent algorithm in GROMACS. The LINCS algorithm was applied to constrain all bond lengths involving hydrogen atoms (*15*). Particle Mesh Ewald was employed to calculate the long-range electrostatic interactions. The cut-off distance for the long-range van der Waals energy term was 10.0 Å. Each system was minimized using the steepest descent algorithm up to 50,000 steps, reducing the maximum force to below 1000 kJ/mol/nm. Temperature equilibration was carried out at 300 K using the V-rescale thermostat for 100 ps with a 2 fs time step (*16*). Pressure equilibration was performed with the C-rescale barostat at 1 bar for 100 ps, also using a 2 fs time step (*17*). Following equilibration, a 5 ns production simulation was conducted. The trajectories were sampled every 10 ps for analysis in production dynamics. RMSD calculations were performed using the ‘gmx rms’ command in GROMACS. RMSF calculations were performed using the ‘gmx rmsf’ command in GROMACS, focusing on the timeframe between 1 ns and 5 ns. The RMSF values were visualised on the protein structures using PyMOL (*18*).

## Acknowledgements

### General

We deeply thank members of Dr. Chikara Frusawa’s lab and Dr. Wataru Iwasaki’s lab for valuable discussions and for helping with computational/experimental setups, especially Mr. Masaki Fujiyoshi, Mr. Kazuki Miyata, Mr. Shun Yamanouchi, Mr. Jun Kuroki, Mr. Keito Watano, and Mr. Yugo Tsunoda. We also appreciate members of Dr. Kozo Tomita’s lab and Dr. Misato Otani’s lab for sharing devices for protein experiments with kind lectures, especially Dr. Yuka Yashiro, Dr. Seisuke Yamashita, and Ms. Natsu Takayanagi. We are deeply indebted to members of Dr. Masahide Kikkawa’s lab for teaching cryo-electron microscopy experiments, especially Ms. Yoriko Akuzawa and Dr. Chieko Saito. We thank Mr. Tatsuki Tabuchi for helping protein experiments. Finally, we express our sincere appreciation to everyone who gave us feedback on this project, especially Mr. Michihiro Nishimura.

### Funding

This work was supported by the Japan Society for the Promotion of Science (KAKENHI Grant numbers 22J20318 and 24H00870 to N.K., JPMJCR19S2 to Y.N., and 21H05247 and 21H05248 to M.K.), the Japan Science and Technology Agency (GteX JPMJGX23B2 to Y.N. and ERATO JPMJER1902 to C. F.), the ANRI Fellowship to N.K., and by Platform Project for Supporting Drug Discovery and Life Science Research (Basis for Supporting Innovative Drug Discovery and Life Science Research (BINDS)) from Japan Agency for Medical Research and Development (AMED) under Grant Number JP24ama121002 to M.K.

### Author contribution

Conceptualization: N.K., Investigation throughout the study: N.K., Docking simulation and molecular dynamics analysis: S.N., Protein expression: N.K. and K.M., Enzymatic assay: N.K., Data visualization: N.K. and S.N., Methodology: N.K., K.M., S.N., K.O., H.Y., S.T., Supervision of cryo-EM analysis: H.Y. and M.K.. Supervision throughout the study: W.I., M.K., and C.F., Writing— original draft: N.K. and S.N., Writing—review and editing: N.K., K.M., S.N., K.O., H.Y., S.T., Y.N., M.K., W.I., and C.F.

### Data availability

The cryo-EM density maps and the atomic coordinates have been deposited in the Electron Microscopy Data Bank (EMDB) and in the PDB, respectively, under the following accession codes: 9LDK (AdhE) and 9 LDL (BdhE) for PDB. EMD-63003 (AdhE) and EMD-63004 (BdhE) for EMDB.

