## Supplementary Figures 1-10 for "Repeatability of protein structural evolution following convergent gene fusions"

**This PDF file includes:**

1. Supplementary Figures 1-10

### Supplementary Figures

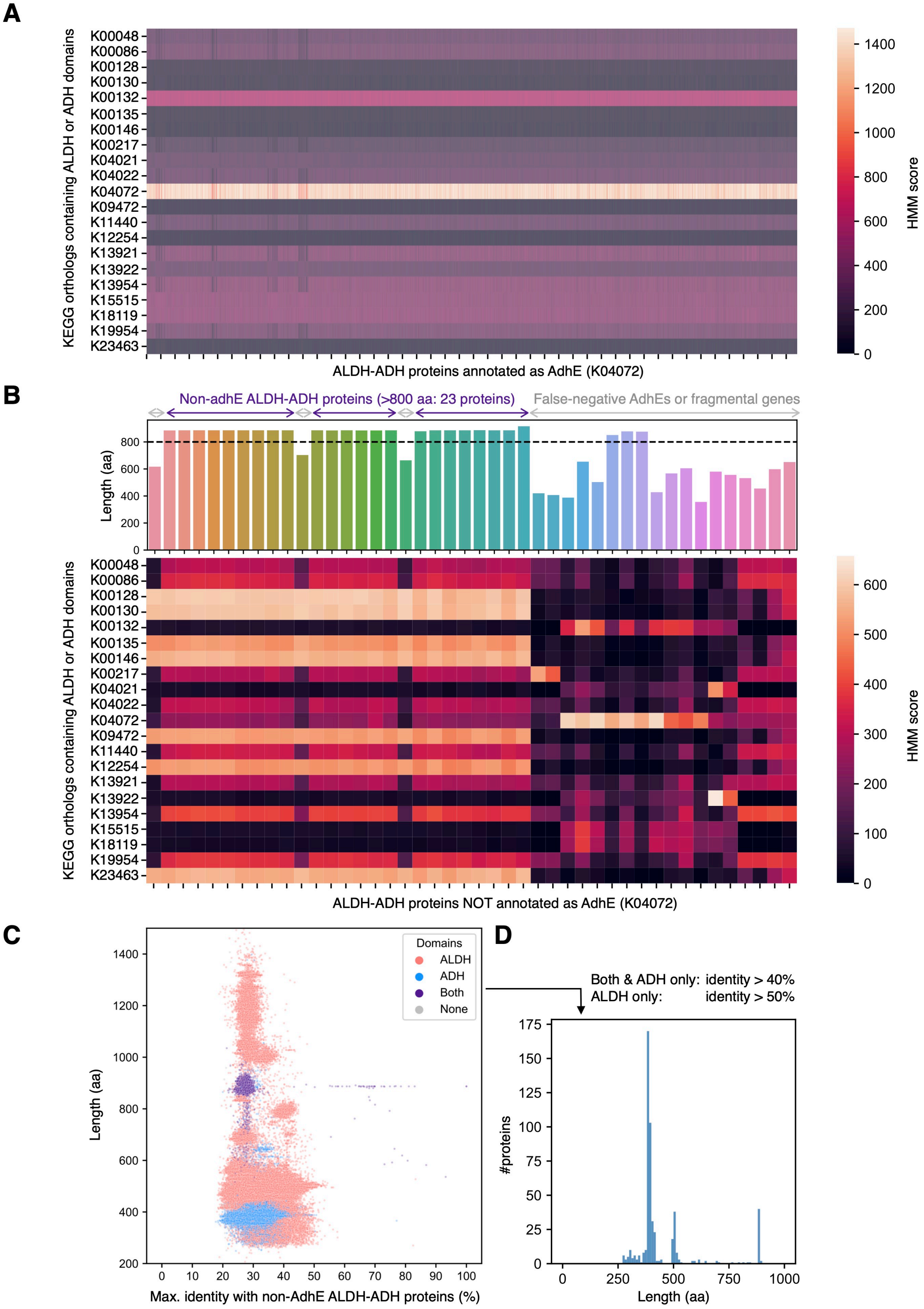

**Supplementary Figure 1. *In silico* extraction of non-AdhE ALDH-ADH fusion proteins. (A)** The results of ortholog annotation for UniProt protein entries annotated as AdhE (K04072) by KofamScan. The heatmap shows Hidden Markov Model (HMM) scores for every AdhE protein and every KEGG Ortholog possessing ALDH or ADH domains. As expected, K04072 showed the highest HMM scores overall. **(B)** The results of ortholog annotation by KofamScan for bacterial proteins NOT annotated as AdhE. The heatmap represents the HMM scores like (A). The bar plot indicates the amino acid length of each protein. As shown in the panel, we focused on the 23 proteins showing >800 aa length and highest HMM scores for non-AdhE orthologs as full-length non-AdhE fusion proteins with ALDH and ADH domains. **(C)** The profiles of the search hit proteins of sequence-similarity-based search by querying the 23 proteins in (B) against all the proteins coded in 45,555 bacterial genomes. Each dot represents each search hit protein, and the X and Y axis represents the maximum alignment identity with query sequences and the amino acid length, respectively. **(D)** The histogram of amino acid length after extracting search hit proteins based on maximum alignment identity. The extraction threshold of alignment identities was 40% and 50% for proteins with both domains or only ADH domain and for those with only ALDH domain, respectively.

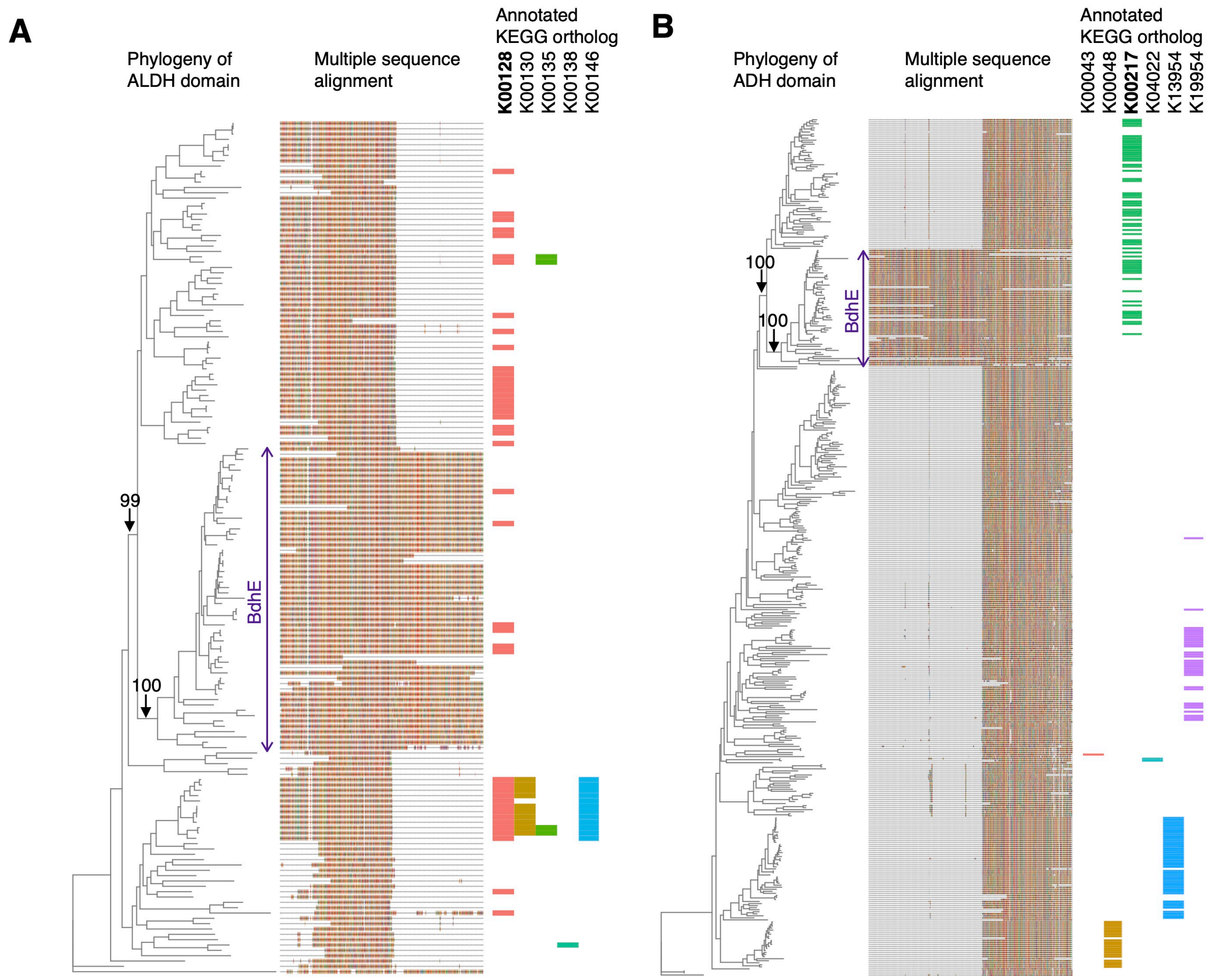


**Supplementary Figure 2. Domain-wise gene phylogenies around non-AdhE ALDH-ADH fusion proteins.** Phylogenetic trees and multiple sequence alignments (MSAs) of the sequence-similarity search hit proteins (**fig. S1D**) with ALDH (A) or ADH domain (B). Ultrafast bootstrap values are shown for specific branches with arrows. Annotated KEGG Orthologs are indicated along with the phylogenies and MSAs.

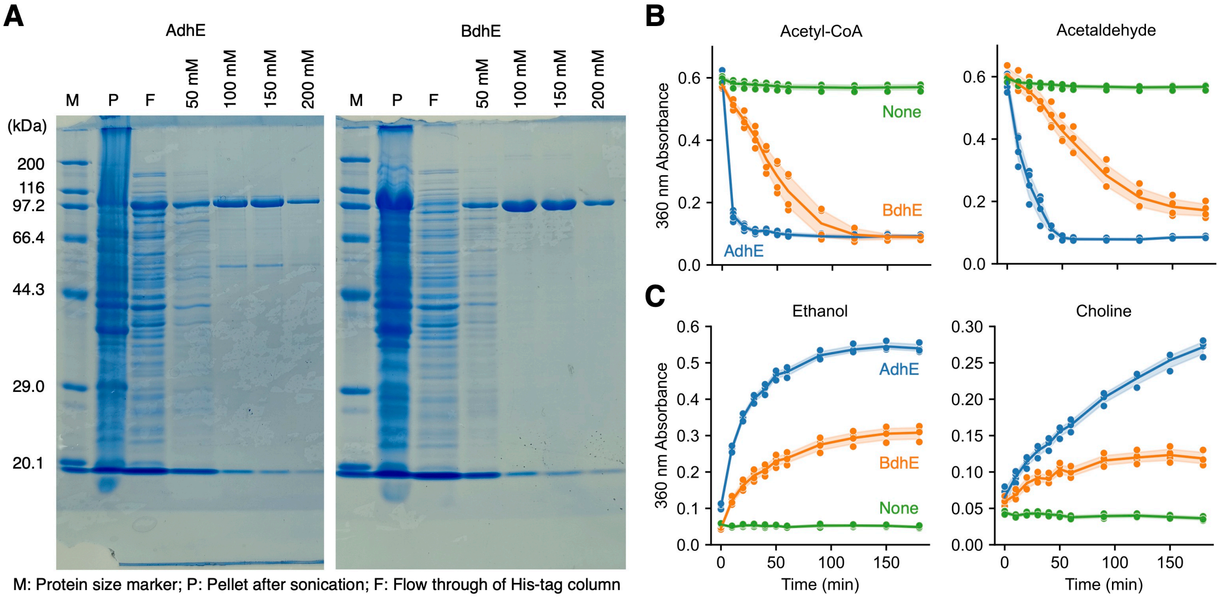


**Supplementary Figure 3. Expression, purification, and enzymatic assays of AdhE and BdhE. (A)** The results of sodium dodecyl sulfate-polyacrylamide gel electrophoresis (SDS-PAGE) for intermediates and final products of expression and His-tag purification of AdhE and BdhE. Lanes “M”, “P”, and “F” indicate molecular weight marker, the centrifugation pellet after sonication, and the flow through of His-tag purification. “50 mM” to “200 mM” indicate the eluate by adding various concentrations (50-200 mM) of imidazole. The molecular weights of AdhE and BdhE monomers with His-tag were 99.7 kDa and 97.9 kDa, respectively. **(B, C)** Enzymatic assays of acetyl-CoA/acetaldehyde oxidization (B) and ethanol/choline oxidization (C) by AdhE and BdhE purified by a His-tag affinity column and a size-exclusion column chromatography. The time-course absorbance of 360 nm was measured as concentrations of NADH for four replicates. “None” represents conditions in which only buffer without enzymes were added.

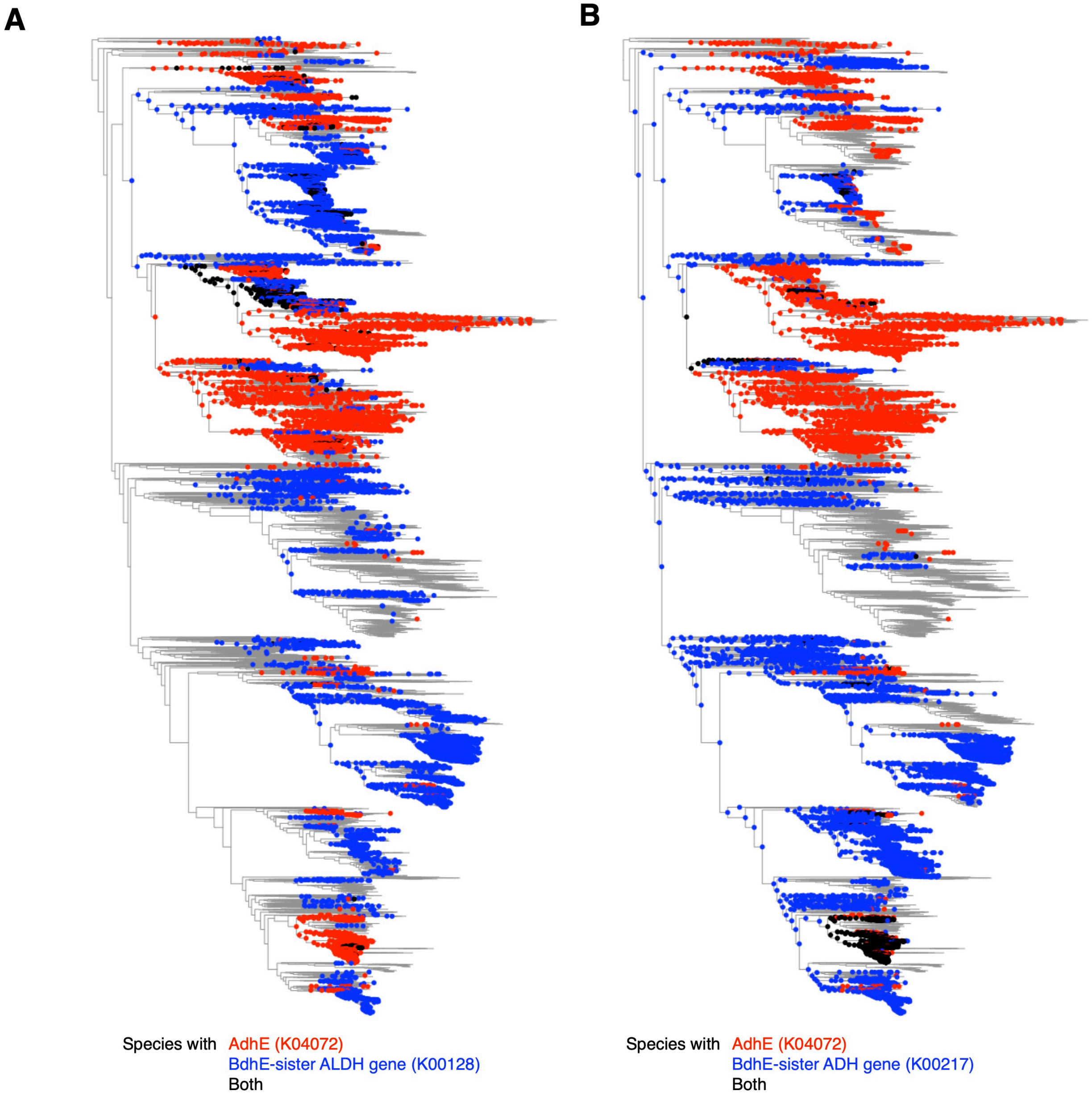


**Supplementary Figure 4. Anti-correlated phylogenetic distributions of *adhE* and *bdhE*-sister genes.** Phylogenetic distribution of *adhE* and *bdhE*-sister genes with ALDH (A) or ADH (B). The phylogenies are bacterial species trees of all the 25,877 species with high-quality genomes (>95% completeness and <5% contamination). The red, blue, and black nodes here indicate the extant/ancestral nodes estimated to possess AdhE, BdhE-sister proteins, and both, respectively. The ancestral states were estimated with PastML v.1.9.15.

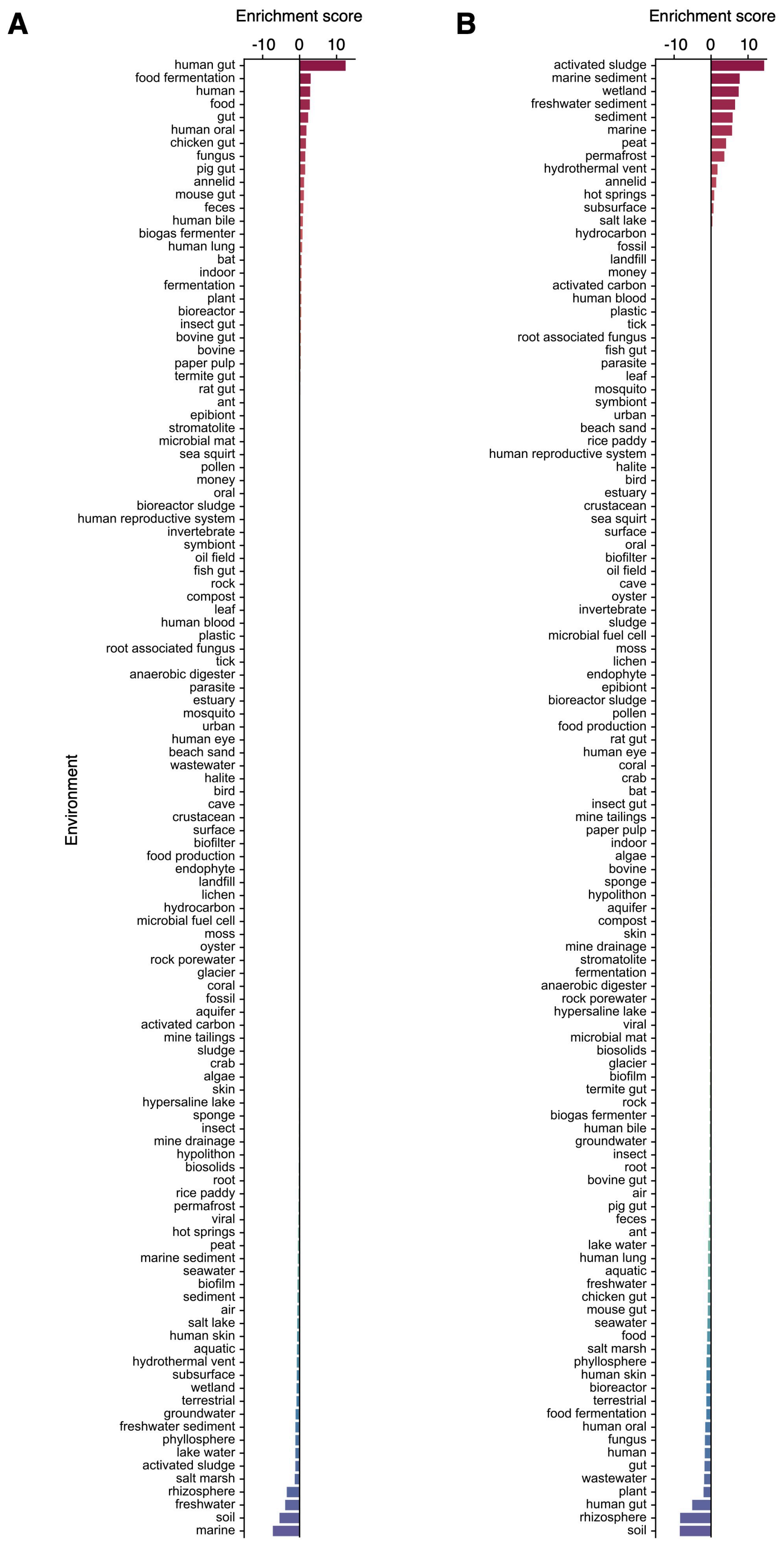
A
Enrichment score
-10
0
10
Environment
human gut
food fermentation
human
food
gut
human oral
chicken gut
fungus
pig gut
annelid
mouse gut
feces
human bile
biogas fermenter
human lung
bat
indoor
fermentation
plant
bioreactor
insect gut
bovine gut
bovine
paper pulp
termite gut
rat gut
ant
epibiont
stromatolite
microbial mat
sea squirt
pollen
money
oral
bioreactor sludge
human reproductive system
invertebrate
symbiont
oil field
fish gut
rock
compost
leaf
human blood
plastic
root associated fungus
tick
anaerobic digester
parasite
estuary
mosquito
urban
human eye
beach sand
wastewater
halite
bird
cave
crustacean
surface
biofilter
food production
endophyte
landfill
lichen
hydrocarbon
microbial fuel cell
moss
oyster
rock porewater
glacier
coral
fossil
aquifer
activated carbon
mine tailings
sludge
crab
algae
skin
hypersaline lake
sponge
insect
mine drainage
hypolithon
biosolids
root
rice paddy
permafrost
viral
hot springs
peat
marine sediment
seawater
biofilm
sediment
air
salt lake
human skin
aquatic
hydrothermal vent
subsurface
wetland
terrestrial
groundwater
freshwater sediment
phyllosphere
lake water
activated sludge
salt marsh
rhizosphere
freshwater
soil
marine
B
Enrichment score
-10
0
10
activated sludge
marine sediment
wetland
freshwater sediment
sediment
marine
peat
permafrost
hydrothermal vent
annelid
hot springs
subsurface
salt lake
hydrocarbon
fossil
landfill
money
activated carbon
human blood
plastic
tick
root associated fungus
fish gut
parasite
leaf
mosquito
symbiont
urban
beach sand
rice paddy
human reproductive system
halite
bird
estuary
crustacean
sea squirt
surface
oral
biofilter
oil field
cave
oyster
invertebrate
sludge
microbial fuel cell
moss
lichen
endophyte
epibiont
bioreactor sludge
pollen
food production
rat gut
human eye
coral
crab
bat
insect gut
mine tailings
paper pulp
indoor
algae
bovine
sponge
hypolithon
aquifer
compost
skin
mine drainage
stromatolite
fermentation
anaerobic digester
rock porewater
hypersaline lake
viral
microbial mat
biosolids
glacier
biofilm
termite gut
rock
biogas fermenter
human bile
groundwater
insect
root
bovine gut
air
pig gut
feces
ant
lake water
human lung
aquatic
freshwater
chicken gut
mouse gut
seawater
food
salt marsh
phyllosphere
human skin
bioreactor
terrestrial
food fermentation
human oral
fungus
human
gut
wastewater
plant
human gut
rhizosphere
soil

**Supplementary Figure 5. Habitat preference of species possessing AdhE or BdhE. (A, B)** The habitat enrichment of species with AdhE or BdhE. The enrichment score represents the difference between each environment's average habitat preference scores calculated for species with and without AdhE (A) or BdhE (B). The habitat preference scores were calculated using the ProkAtlas pipeline (see MATERIALS AND METHODS).

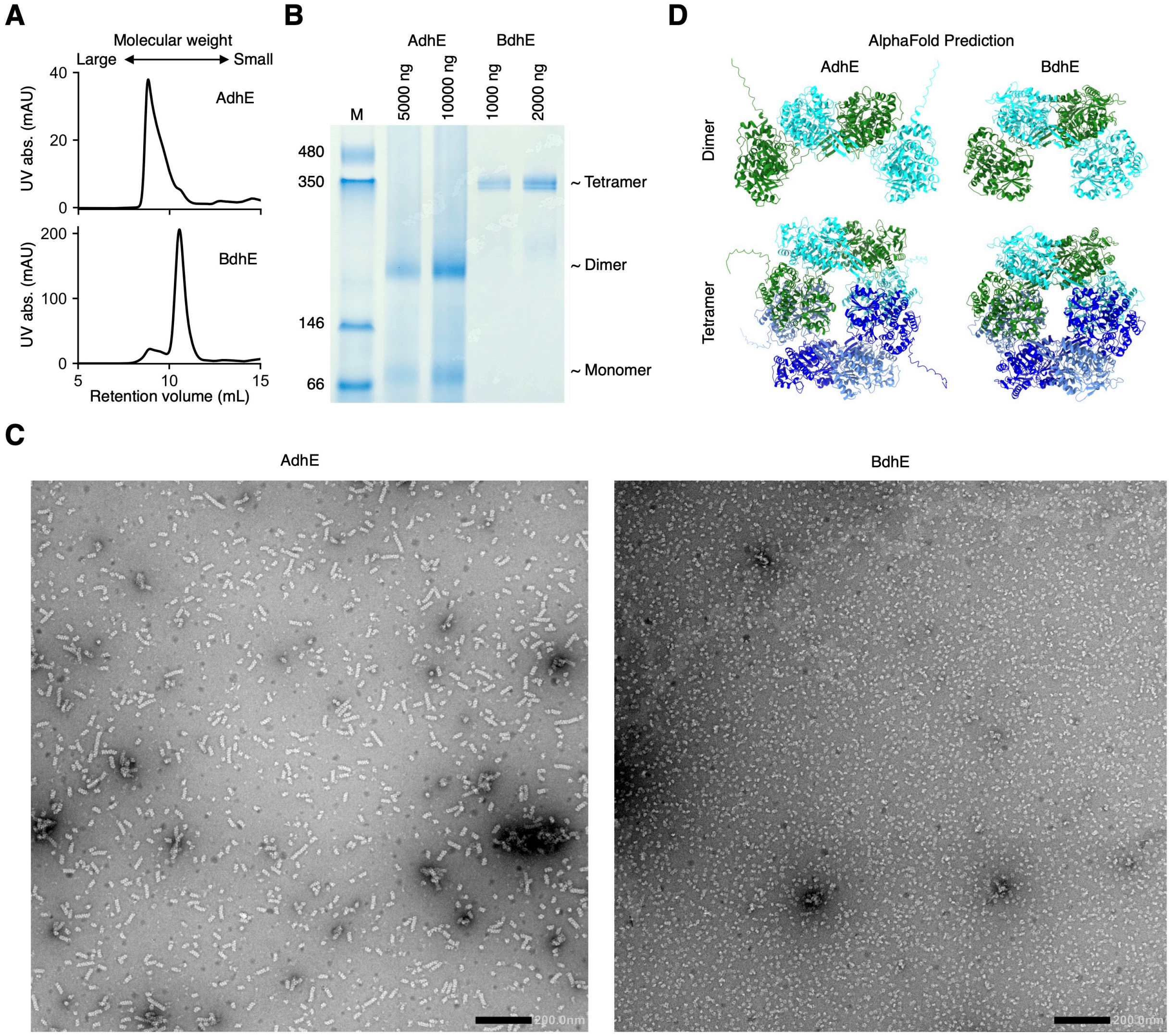


**Supplementary Figure 6. Multimer stoichiometry and structural analysis for AdhE and BdhE. (A)** Chromatograms of AdhE and BdhE by gel filtration chromatography after his-tag purification. **(B)** Blue native PAGE for AdhE and BdhE after his-tag purification. The applied weights of proteins in loaded samples are indicated above the gel. **(C)** The full negative stain electron microscopy analysis of AdhE and BdhE. Scale bar 200.0 nm. **(D)** Dimer and tetramer structures of AdhE and BdhE from *E. coli* and *H. eurihalina* predicted by AlphaFold v2.3.2. Different molecules are colored differently.

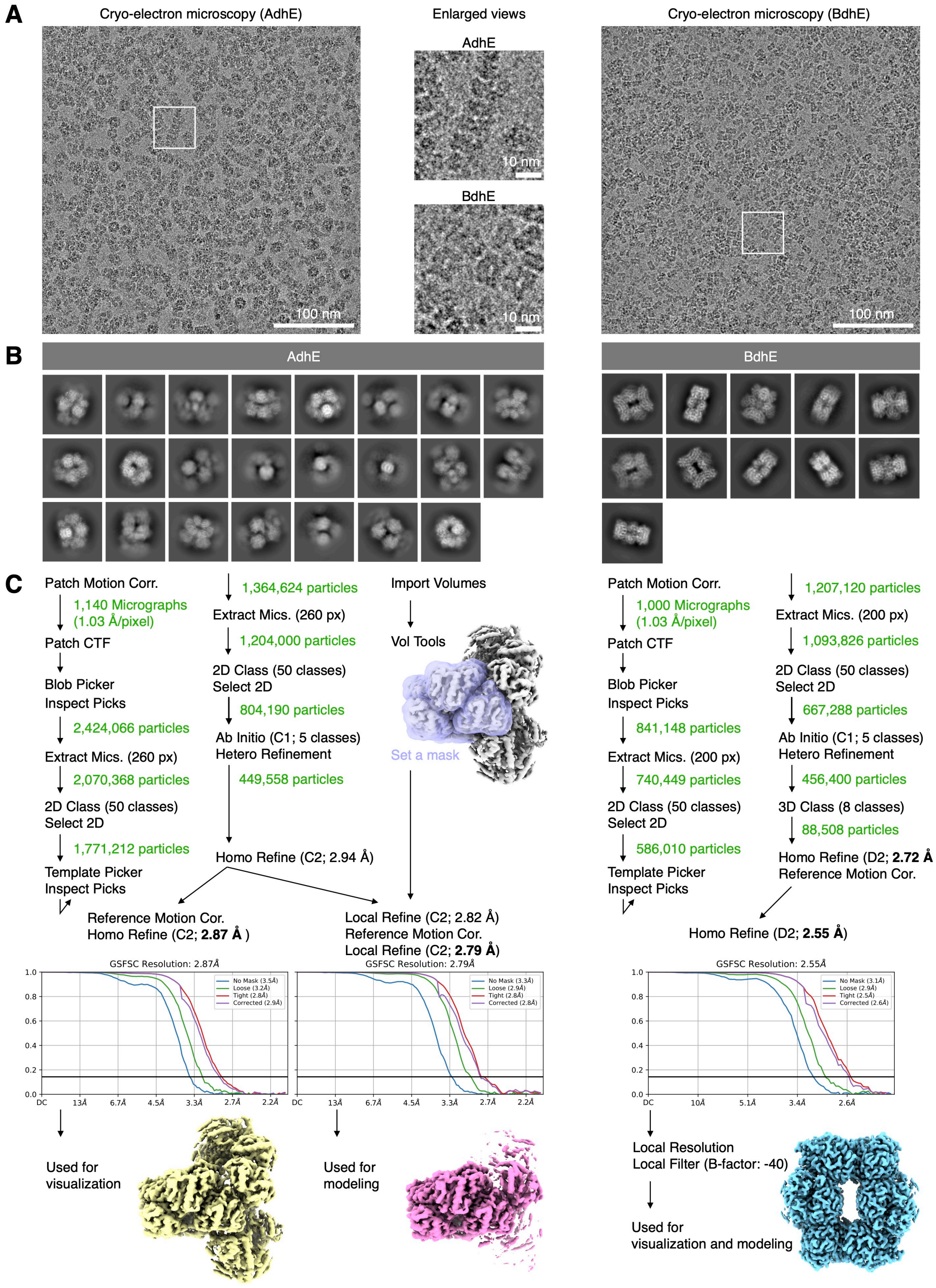
A
Cryo-electron microscopy (AdhE)
Enlarged views
Cryo-electron microscopy (BdhE)
AdhE
BdhE
100 nm
10 nm
10 nm
100 nm
B
AdhE
BdhE
C
Patch Motion Corr.
1,140 Micrographs
(1.03 Å/pixel)
Patch CTF
Blob Picker
Inspect Picks
2,424,066 particles
Extract Mics. (260 px)
2,070,368 particles
2D Class (50 classes)
Select 2D
1,771,212 particles
Template Picker
Inspect Picks
1,364,624 particles
Extract Mics. (260 px)
1,204,000 particles
2D Class (50 classes)
Select 2D
804,190 particles
Ab Initio (C1; 5 classes)
Hetero Refinement
449,558 particles
Homo Refine (C2; 2.94 Å)
Import Volumes
Vol Tools
Set a mask
Reference Motion Cor.
Homo Refine (C2; 2.87 Å )
Local Refine (C2; 2.82 Å)
Reference Motion Cor.
Local Refine (C2; 2.79 Å)
Patch Motion Corr.
1,000 Micrographs
(1.03 Å/pixel)
Patch CTF
Blob Picker
Inspect Picks
841,148 particles
Extract Mics. (200 px)
740,449 particles
2D Class (50 classes)
Select 2D
586,010 particles
Template Picker
Inspect Picks
1,207,120 particles
Extract Mics. (200 px)
1,093,826 particles
2D Class (50 classes)
Select 2D
667,288 particles
Ab Initio (C1; 5 classes)
Hetero Refinement
456,400 particles
3D Class (8 classes)
88,508 particles
Homo Refine (D2; 2.72 Å
Reference Motion Cor.
Homo Refine (D2; 2.55 Å)
GSFSC Resolution: 2.87Å
No Mask (3.5Å)
Loose (3.2Å)
Tight (2.8Å)
Corrected (2.9Å)
GSFSC Resolution: 2.79Å
No Mask (3.3Å)
Loose (2.9Å)
Tight (2.8Å)
Corrected (2.8Å)
GSFSC Resolution: 2.55Å
No Mask (3.1Å)
Loose (2.9Å)
Tight (2.5Å)
Corrected (2.6Å)
Used for
visualization
Used for
modeling
Local Resolution
Local Filter (B-factor: -40)
Used for
visualization and modeling

**Supplementary Figure 7. Workflows of cryo-electron microscopy analysis. (A)** An example micrograph of cryo-electron microscopy for AdhE and BdhE. The enlarged view of each micrograph is shown in the middle, and the enlarged regions are shown with white boxes in the full original micrographs. In comparison with the negative stain image (Fig. S6C), a larger proportion of AdhE particles appeared as top views rather than side views of filaments. This could be due to the random orientation of short AdhE filaments in the frozen sample for the cryo-electron microscopy experiments. **(B)** Reconstructed 2D images of AdhE and BdhE after 2D classification of particles. Each image corresponds to each particle class. **(C)** Pipelines to reconstruct electron density maps of AdhE and BdhE. CTF refinement was conducted in the homogeneous refinement process (Homo Refine).

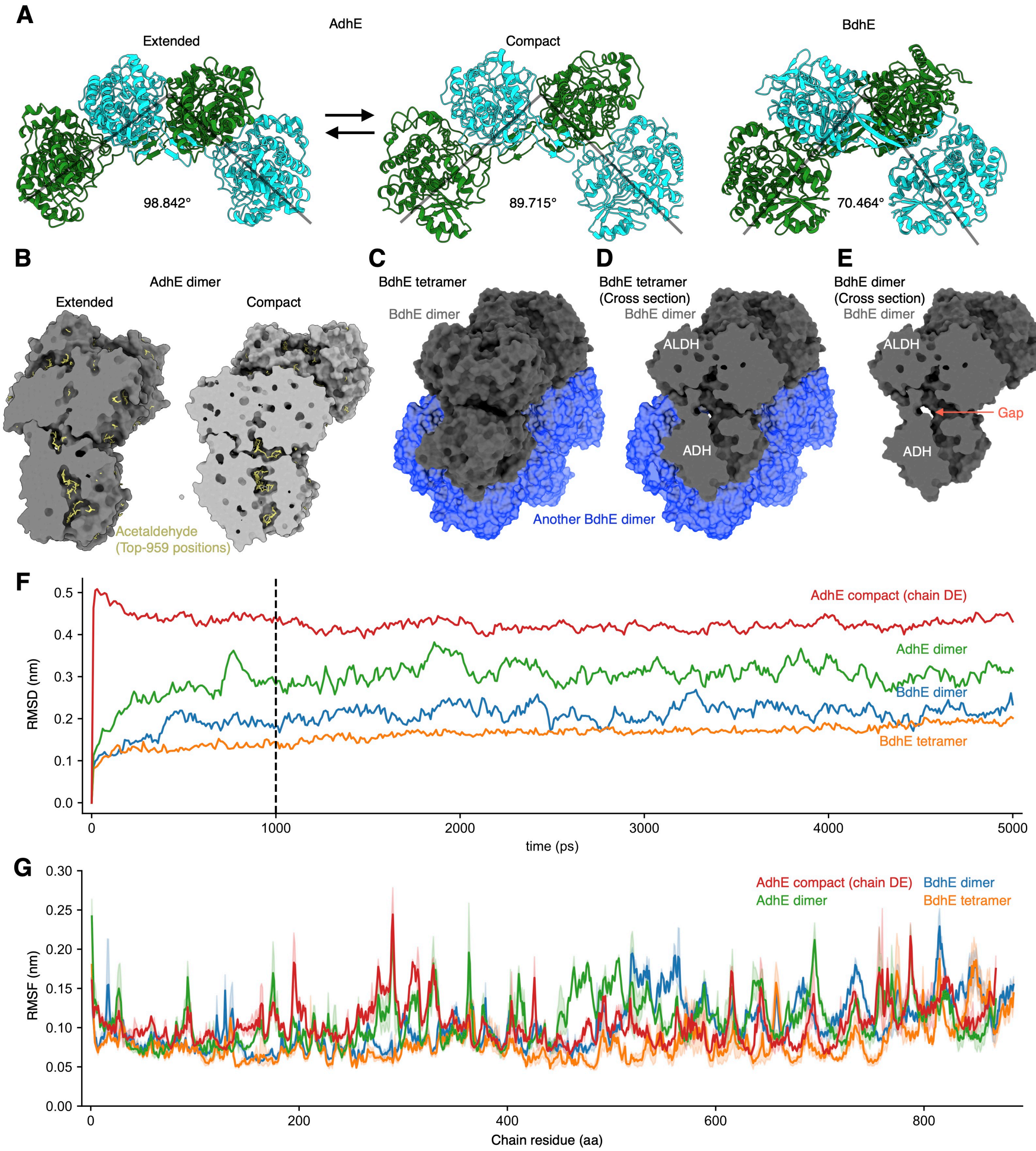


**Supplementary Figure 8. Comparison of multimeric structures and their stability of AdhE and BdhE. (A)** The dimeric structure units of AdhE and BdhE. The bending angles of AdhE and BdhE represent angles formed by connecting C-α atoms at L638 and P208 of a chain, and L638 of the other chain in AdhE, F681 and K250 of a chain, and F681 of the other chain in BdhE, respectively. For AdhE, the angle was calculated for two distinct conformations (extended (PDB ID: 6TQH) and compact (PDB ID: 6TQM)). **(B)** Docking simulation results of AdhE dimers and an acetaldehyde. Docking simulations were conducted for AdhE dimers of extended and compact conformation. The yellow molecular structures represent 959 possible docking positions of acetaldehyde within 4 kcal/mol from the lowest-energy position. **(C-E)** Gap filling of substrate tunnel wall in a BdhE dimer by another dimer in BdhE tetramers**.** The surface structure of BdhE tetramer (C), the cross-section of the tetramer (D), and a cross-section of single dimeric structure unit of BdhE (E). The structural gap of the substrate tunnel is indicated with an orange allow (E), but is partially filled by binding with another dimeric unit (D). **(F)** Time-course tracking of RMSD for AdhE and BdhE's structures while molecular dynamics simulations and their initial structures. The results for AdhE hexamer/dimer and BdhE tetramer/dimer are shown. **(G)** RMSFs calculated for every residue. RMSFs in multiple chains were averaged.

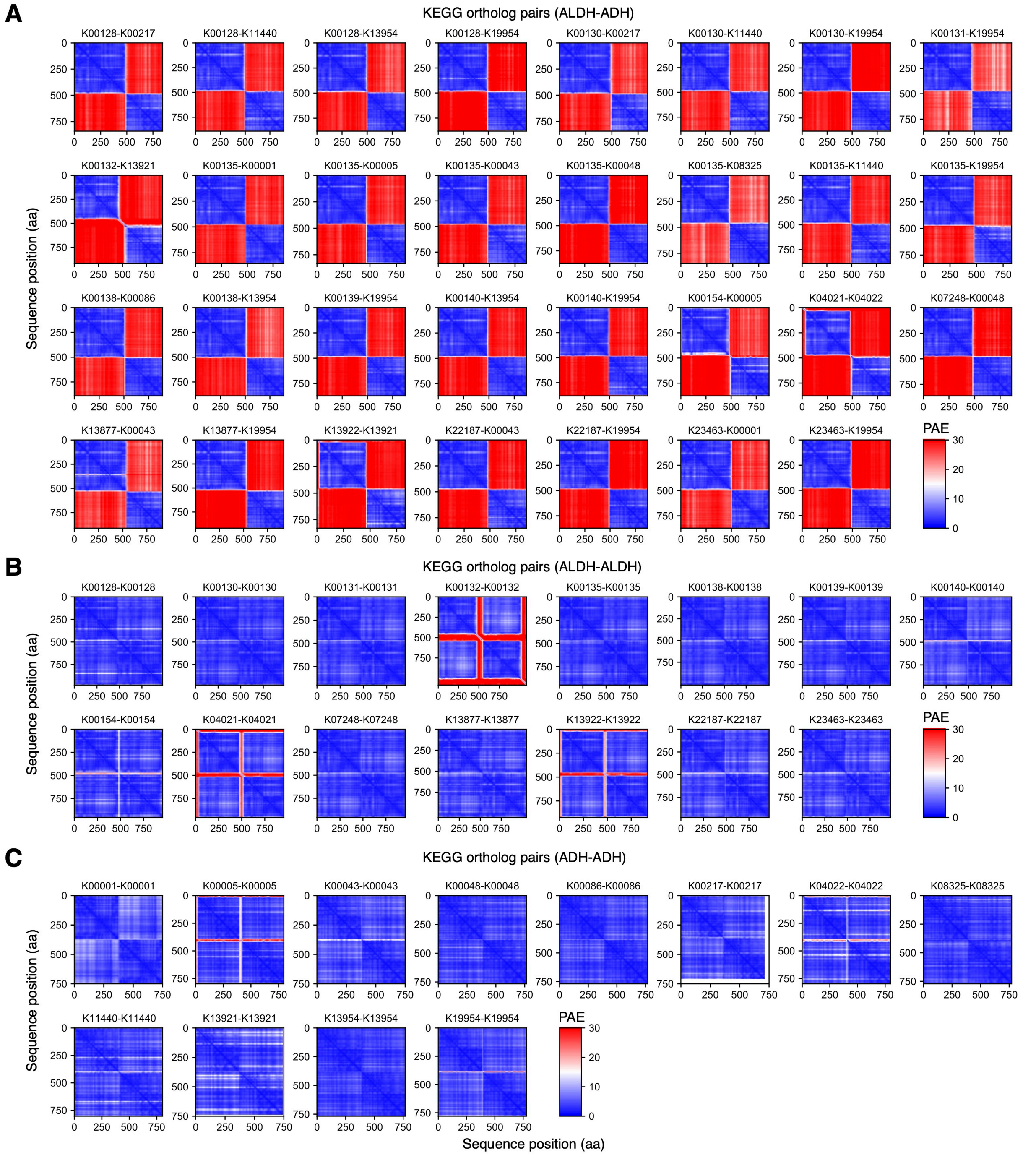


**Supplementary Figure 9. ColabFold-predicted heteromeric and homomeric interactions of ALDH and ADH proteins. (A)** ColabFold structure prediction of heteromeric interaction between ALDH and ADH genes adjacently coded in genomes. Structural predictions were done by ColabFold v1.5.2. The heatmap indicates the positioned aligned errors (PAEs). **(B, C)** ColabFold structure prediction of homodimeric interactions, i.e., ALDH-ALDH (B), and ADH-ADH (C) contacts.

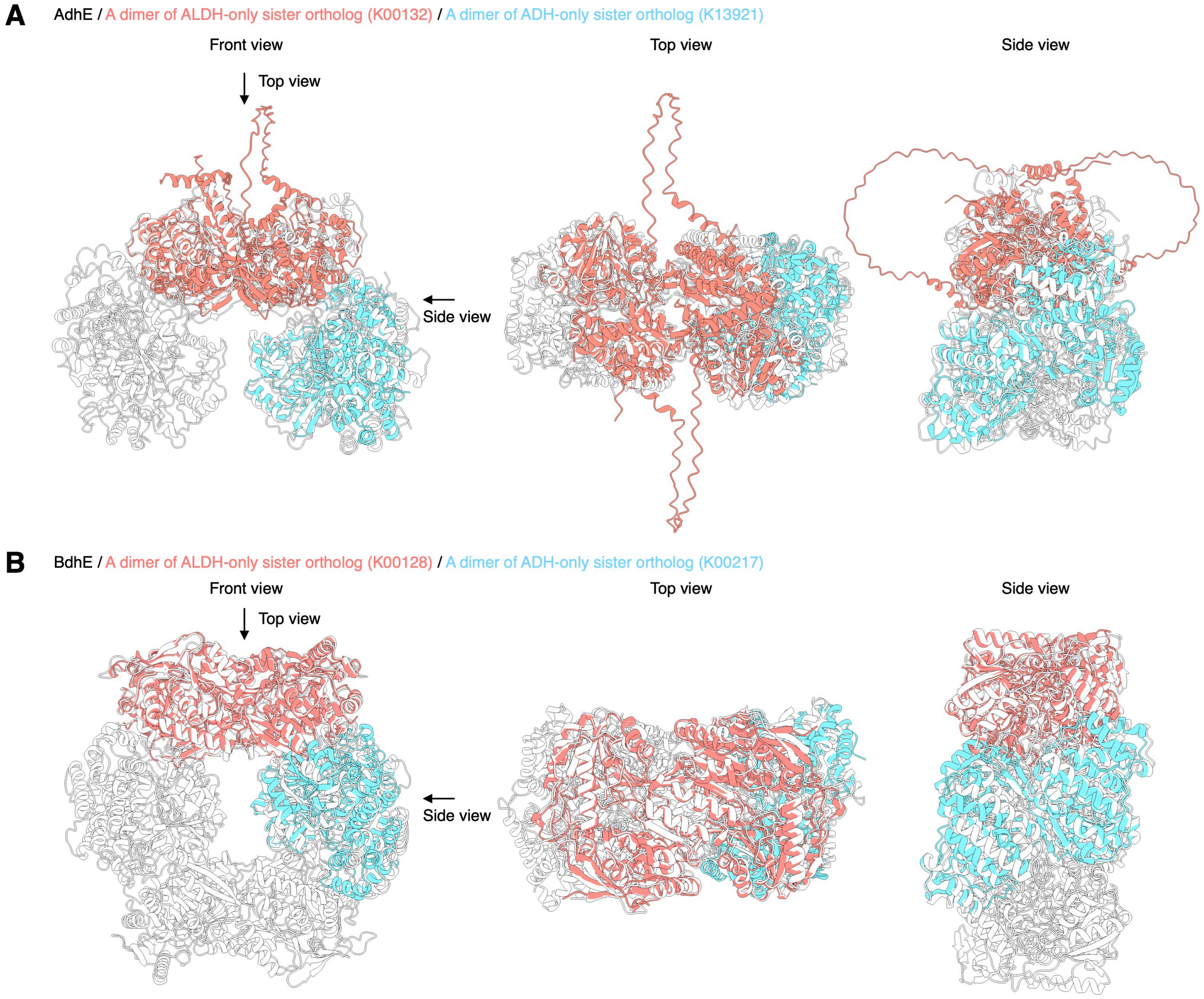


**Supplementary Figure 10. Structural alignment of AdhE/BdhE and the predicted structures of the sister-clade proteins. (A, B)** Three-dimensional structure of AdhE (A; PDB ID: 6TQM) or BdhE (B), aligned with a ColabFold-predicted dimeric structure of a randomly selected protein from the sister clade of AdhE or BdhE.
